# HNRNPM controls circRNA biogenesis and splicing fidelity to sustain prostate cancer cell fitness

**DOI:** 10.1101/2020.06.17.157537

**Authors:** Jessica SY Ho, Diana Low, Megan Schwarz, Danny Incarnato, Florence Gay, Tommaso Tabaglio, Jingxian Zhang, Heike Wollman, Leilei Chen, Omer An, Tim Hon Man Chan, Alexander Hall Hickman, Simin Zheng, Vladimir Roudko, Sujun Chen, Musaddeque Ahmed, Housheng Hansen He, Benjamin D. Greenbaum, Ivan Marazzi, Michela Serresi, Gaetano Gargiulo, Salvatore Oliviero, Dave Keng Boon Wee, Ernesto Guccione

**Affiliations:** Institute of Molecular and Cell Biology (IMCB), Agency for Science, Technology and Research (A*STAR), Singapore; Department of Biochemistry, Yong Loo Lin School of Medicine, NUS, Singapore; Cancer Science Institute of Singapore, National University of Singapore, Singapore 117599, Singapore; Center for Therapeutics Discovery, department of Oncological Sciences and Pharmacological Sciences, Tisch Cancer Institute, Icahn School of Medicine at Mount Sinai, New York, New York, USA; IIGM (Italian Institute for Genomic Medicine), Torino Italy; Dipartimento di Scienze della Vita e Biologia dei Sistemi Università di Torino Via Accademia Albertina 13 Torino Italy., Torino, Italy; University of Groningen; Department of Anatomy, Yong Loo Lin School of Medicine, National University of Singapore, Singapore 117594, Singapore; School of Biological Sciences, Nanyang Technological University, Singapore, Singapore; Tisch Cancer Institute, Icahn School of Medicine at Mount Sinai, New York, NY 10029, USA; Department of Medicine, Hematology and Medical Oncology, Icahn School of Medicine at Mount Sinai, New York, NY 10029, USA; Department of Oncological Sciences, Icahn School of Medicine at Mount Sinai, New York, NY 10029, USA; Department of Pathology, Icahn School of Medicine at Mount Sinai, New York, NY 10029, USA; Department of Medical Biophysics, University of Toronto, Toronto, Ontario, Canada; Princess Margaret Cancer Center, University Health Network, Toronto, Ontario, Canada; Ontario Institute for Cancer Research, Toronto, Ontario, Canada; Max Delbruck Center for Molecular Medicine, Berlin-Buch, Germany; The Netherlands Cancer Institute, Amsterdam, The Netherlands; Department of Microbiology, Icahn School of Medicine at Mount Sinai, New York. USA

**Author notes:** Correspondence should be addressed to Ernesto Guccione.

**Keywords:** hnRNPM, Prostate Cancer, splicing, circular RNA, Chromatin, histone methylation, EED, H3K27me3

## Abstract

Cancer cells are differentially dependent on the splicing machinery compared to normal untransformed cells. The splicing machinery thus represents a potential therapeutic target in cancer. To identify splicing factors important for prostate cancer cell (PCa) cell growth, we performed a parallel pooled shRNA screen on *in vitro* passaged cells and *in vivo* xenografted PCa tumor lines. Our screen revealed HNRNPM as a potential regulator of PCa cell growth. RNA- and eCLIP-sequencing data suggest that HNRNPM is bound to transcripts of key homeostatic genes and that loss of HNRNPM binding in a subset of these genes results in aberrant exon inclusion and exon back-splicing events in target transcripts. In both linear and circular mis-spliced transcripts, HNRNPM appears to preferentially bind to GU-rich elements in long flanking proximal introns. Mimicry of HNRNPM dependent linear splicing events using splice-switching antisense oligonucleotides (SSOs) was sufficient to inhibit cell growth in HNRNPM expressing cells. This suggests that prostate cancer cell dependence on HNRNPM is likely a result of mis-splicing of key homeostatic coding and non-coding genes. Taken together, our data reveal a role for HNRNPM in supporting prostate cancer cell fitness, and also as a potential therapeutic target in PCa.

## INTRODUCTION

In eukaryotic cells, many genes harbor intronic sequences that are removed during RNA splicing and transcript maturation (Venter et al., 2001). This process is regulated by the spliceosome, comprising of small non-coding RNAs (U1, U2, U4, U5 and U6), the core spliceosomal proteins (U2AF1, U2AF2, SF3B1 etc.) and other auxiliary factors (Wahl, Will, & Luhrmann, 2009). Under normal physiological conditions, proper regulation of splicing provides the cell an opportunity to control gene expression in the absence of genetic alterations. By expressing alternative isoforms of the same gene, the cell can regulate both the inclusion/exclusion of specific protein and/or RNA domains (Ellis et al., 2012; Yang et al., 2016) or change transcript half-life (Lareau & Brenner, 2015; Naro et al., 2017; t Hoen et al., 2011), thus impacting both transcript fate and function.

The regulation of alternative splicing plays a central role in development (Baralle & Giudice, 2017), cellular differentiation (Pimentel et al., 2016) as well as in the cellular response to external or internal stimuli (Hang et al., 2009; Haque, Ouda, Chen, Ozato, & Hogg, 2018; Makino, Kanopka, Wilson, Tanaka, & Poellinger, 2002; Shalgi, Hurt, Lindquist, & Burge, 2014). However, the same phenotypic plasticity offered by the splicing machinery can work against the cell and organism, and confer competitive advantages to cells under pathological conditions such as cancer. Indeed, alterations in the expression of specific isoforms of certain genes and of splicing factors themselves can promote cell proliferation (e.g. Androgen Receptor (AR); Liu et al. (2014)), metastasis (e.g. CD44; (Todaro et al., 2014; Xu et al., 2014)) or avoid apoptosis (Dewaele et al., 2016; Schwerk & Schulze-Osthoff, 2005) in cancer. Some of these changes in isoform ratios may be direct results of mutations in the linear motifs that comprise the splice sites (Bartram et al., 2017; Jung et al., 2015; Puente et al., 2015) or in the splicing machinery itself (Graubert et al., 2011; Papaemmanuil et al., 2011; Wang et al., 2011; Yoshida et al., 2011).

In addition to the differential splicing of specific genes, there is also increasing evidence that some molecular subtypes of cancer, which bear mutations in non-splicing related genes such as MYC, are highly dependent on a functional core spliceosome for survival (Hsu et al., 2015; Koh et al., 2015). This may be related to their high proliferation rates, which require them to heavily rely on the splicing machinery.

Thus, there is evidence for a strong involvement of the RNA splicing in cancer. The splicing machinery and its associated factors may offer selective therapeutic vulnerabilities. However, the identify of critical targets, the context in which they function, their mechanisms of action, and the functional impact of the individual splicing factors in cancer remain unknown. To address this, we performed a parallel pooled *in vivo* and *in vitro* shRNA screens against core and auxiliary splicing factors in prostate cancer (PCa). Through this screen, we were able to uncover a novel role for the heterogeneous nuclear ribonucleoprotein M (HNRNPM). HNRNPM expression is higher in PCa than in normal prostate epithelial cells, and its loss affects PCa cell growth *in vitro* and *in vivo*. RNA-seq and eCLIP suggests that HNRNPM binding differentially impacts the splicing and/or expression of distinct classes of transcripts. Interestingly, we found that HNRNPM is limits circular RNA biogenesis within cells. Overall, our work offers mechanistic insight into the function of HNRNPM in prostate cancer and suggests that HNRNPM could serve as a potential marker or drug target against prostate cancer.

## RESULTS

### A targeted pooled shRNA screen identifies splicing-related dependencies in PCa

In order to identify major splicing factor dependencies in prostate cancer, we conducted parallel pooled *in vitro* and *in vivo* shRNA screens in the LNCAP prostate cancer cell line (**Fig 1A**). In such a screen, shRNAs targeting essential genes and oncogenes are generally depleted as the cell population increases, whereas shRNAs that target tumor suppressors and negative regulators of proliferation become enriched. The screens utilized a lentiviral library of 520 shRNAs targeting 102 core and auxiliary splicing factors, meaning 3 to 5 shRNAs targeting each gene (**Table S1**). After puromycin selection, we harvested a portion of the cells to be used as the control input pool (P1). The remaining cells were either passaged continuously *in vitro* for about 28-30 population doublings (about 1 month; ~9 passages), or xenografted into the flanks of SCID mice and allowed to form tumors over time. *In vitro* passaged cells were harvested at every other passage for DNA isolation (P3, P5, P7, P9). *In vivo* tumors were harvested for DNA isolation when they attained a size of 400mm^3^ or more. shRNA hairpin enrichment levels in individual samples were quantified via high throughput sequencing. Input lentiviral and plasmid pools were also sequenced as controls.

**Figure 1:**
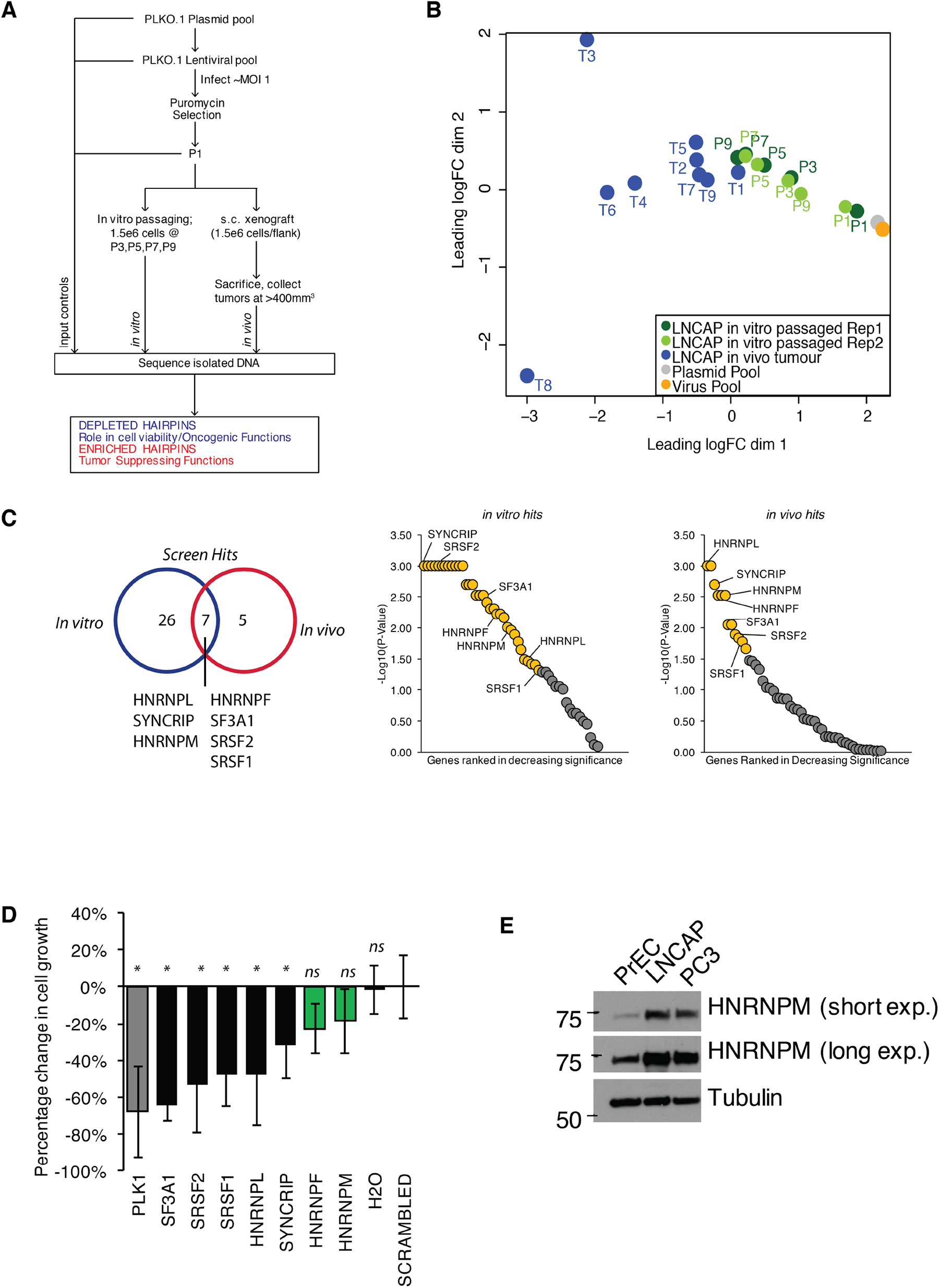
A pooled shRNA screen identifies HNRNPM as a regulator of PCa cell growth. **(A)** Schematic of overall experimental design. **(B)** MDS plot of individual samples (*in vitro* passaged cells and tumors) collected in the screen, in relation to each other. **(C)** (Right) Venn Diagram showing overlap of significant hits for *in vitro* and *in vivo* screens (Left) Ranked dotplots showing −log10(p-values) of *in vivo* and *in vitro* screen hits. Red circles represent significant hits. Tumor-specific hits are indicated **(D)** Barplots showing overall PrEC cell proliferation upon a 96 hour, siRNA mediated knockdown of the indicated splicing factors, as measured through a colorimetric MTS assay. siRNAs that inhibit cell proliferation relative to the scrambled siRNA control are indicated in black, while siRNAs that do not alter cell proliferation are indicated in green. Each assay was performed in 2 biological replicates, with two technical duplicates per replicate. Error bars represent s.d, *: p<0.05 compared to scrambled. **(E)** Western blot of HNRNPM protein levels in PrEC, LNCAP and PC3 cells. Tubulin is shown as a loading control.

For analysis, individual hairpin abundances were normalized to the control input pool (P1). Multidimensional scaling (MDS) analysis of the data showed P1 clustering with the plasmid and lentiviral controls, confirming that hairpin abundances in our input pool of cells were representative of our initial library (**Fig 1B**). Tumor samples were clustered away from P1 samples, indicating a shift in overall hairpin distribution in these samples. Notably, for the *in vitro* passaged samples, samples derived from the later passages clustered closer to tumor samples, while samples derived from the earlier passages clustered more closely to the input group of samples (**Fig 1B**), possibly underscoring similarities in the functional enrichment/depletion patterns during cell proliferation. Consistently, we identified five well-defined clusters of hairpin enrichment/depletion, with similar patterns of enrichment/depletion in both *in vitro* passaged and *in vivo* tumor samples (**Fig S1A**).

To identify top hits in our screen, we subjected our dataset to a rotational gene set analysis (Wu et al., 2010) (**Fig 1C; Tables S2 and S3**). This analysis takes into account the overall enrichment levels of individual hairpins targeting the same gene and compares them against each other. Because the majority of hairpins in our screen appeared to be depleted over time both *in vitro* and *in vivo* (**Fig S1A**), this suggested that many of the hits we found were likely to be essential genes within cells. To distinguish between essential genes from genes that had a potential oncogenic function, we further performed a short-term (96 hours), independent siRNA screen (**Fig 1D**) for our top hits (two-sided p ≤ 0.01, FDR≤0.05) on normal, untransformed prostate epithelial cells (PrEC). We rationalized that an acute reduction in expression of essential genes would likely be deleterious in these cells. On the other hand, reduced expression of genes that were more likely to be important for oncogenic growth should minimally inhibit PrEC cell growth. Indeed, short term, reduced expression of several genes, such as *SF3A1, SRSF1, SRSF2*, which are essential genes that form part of the core spliceosomal machinery, was rapidly deleterious in the PrEC cells (**Fig 1D**). On the other hand, reduced expression of *HNRNPM* and *HNRNPF* mildly affected proliferation in the PrEC cell line, suggesting a long-term oncogenic as opposed to an essential role for these genes in PCa (**Fig 1D**). To further narrow down the hits from our screen, we reanalyzed publicly available gene expression data sets for prostate cancer (TGCA). This analysis correlated the increased expression of *HNRNPM* with poorer disease-free survival over time, while this was not the case for *HNRNPF* (**Fig S1B**). In addition, western blot analysis suggested that expression of *HNRNPM* is increased in PCa cells, but not in normal PrEC cells (**Fig 1E**). Taken together, *HNRNPM* is a valid target for further analysis.

### HNRNPM is required for PCa cell proliferation *in vitro* and *in vivo*

We first validated our screen results using shRNAs (TRCN0000001244; 2B7, TRCN0000001246; 2B9) present in the screen (**Fig 2A and 2B**). These hairpins were efficient in reducing HNRNPM protein and RNA levels (**Fig 2A and 2B**). In line with the screen results, reduced *HNRNPM* mRNA and protein levels in LNCAP cells resulted in reduced cell proliferation (**Fig 2C**). Moreover, we showt that HNRNPM depletion imparirs colony formation (**Fig 2D**) and anchorage-independent cell growth (**Fig 2E**), underscoring a broad role for *HNRNPM* in preserving cell fitness. *HNRNPM* dependency appeared similar in the PC3 prostate cancer cell line (**Fig S2A-E**), which bears different driver mutations (*PTEN* null, *TP53* null, Androgen independent) from LNCAP (*PTEN* null, *TP53* wildtype, Androgen dependent)(van Bokhoven et al., 2003). This suggests that the impact of HNRNPM in PCa is not dependent on specific tumor genotypes or hormone-dependency. Finally, the ability of LNCAP cells to form tumors when xenografted in mice was impaired in HNRNPM-depleted as compared to control cells (**Fig 2F, Fig S2F**). The reduction in cell proliferation and growth correlated with potency of HNRNPM protein depletion by individual hairpins, supporting an on-target effect (**Fig 2 and Fig S2**). These results suggest that HNRNPM is important for maintaining PCa cell growth both *in vitro* and *in vivo*.

**Figure 2:**
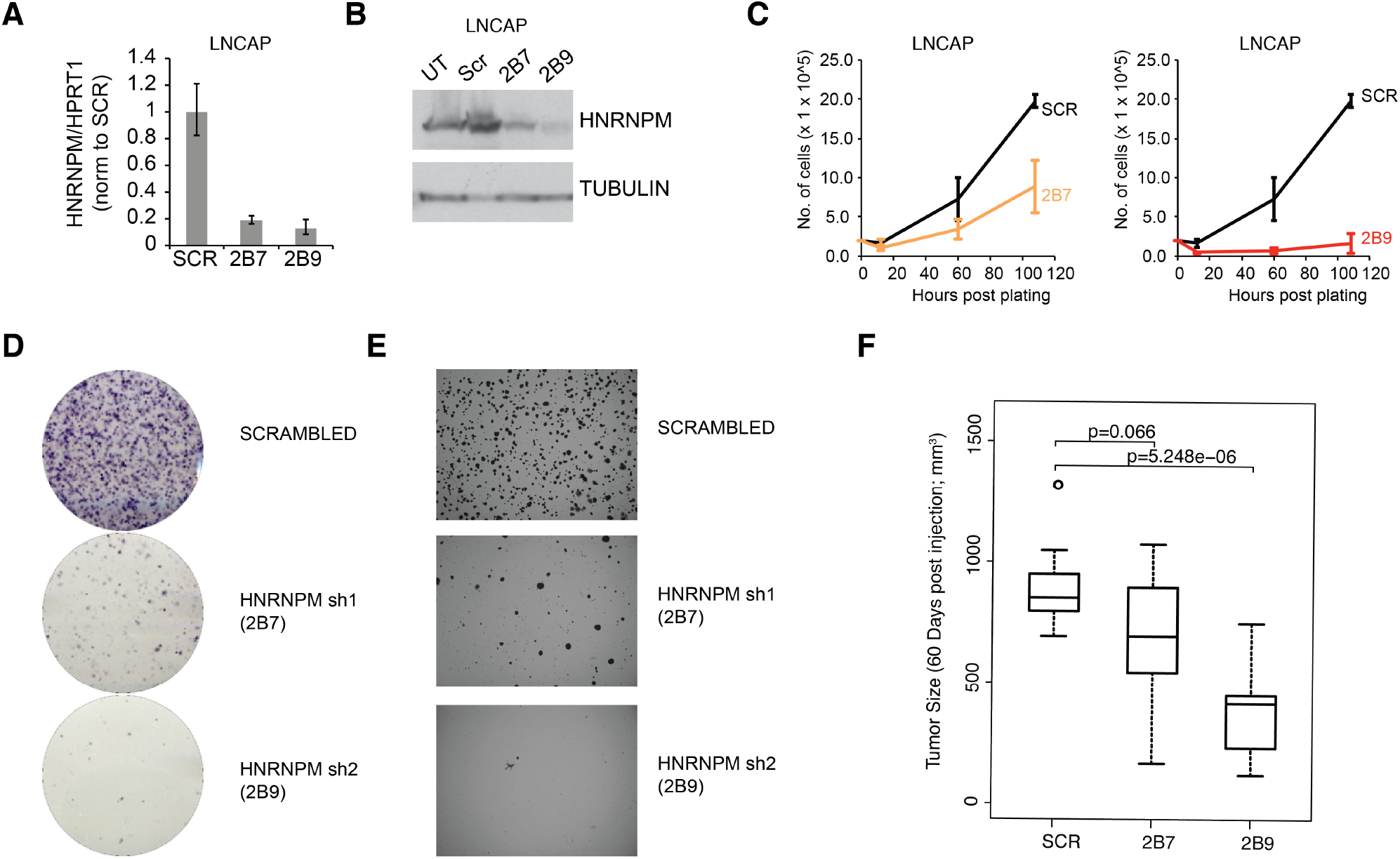
HNRNPM inhibits PCa cell growth in vitro and in vivo. HNRNPM RNA **(A)** and protein levels **(B)** upon expression of scrambled or HNRNPM specific shRNAs in LNCAP cells. **(C)** Cell proliferation assays of LNCAP cells expressing with either scrambled or HNRNPM specific shRNAs (2B7 and 2B9). **(D)** colony formation assays and **(E)** Anchorage independent growth (soft agar assays) of shRNA treated LNCAP cells **(F)** Size of xenografted, scrambled or HNRNPM specific shRNA expressing LNCAP cells at 60 days post injection in 6-8 week old SCID-CB17 mice (n=10-11 per condition).

### HNRNPM is bound to GU-rich elements within long genes

To further understand the molecular role of HNRNPM in PCa, we next performed eCLIP (Van Nostrand et al., 2016) on LNCAP cells to identify direct RNA targets of HNRNPM. eCLIP analysis (**Fig S3A-C**; eCLIP controls) revealed 57984 high confidence (p-value <= 0.05, Fold enrichment >= 1.5) HNRNPM bound sites in LNCAP cells (**Fig 3A, S3B and S3C**), spanning 6510 gene bodies. These sites were primarily located in intronic regions of expressed genes, with very few binding sites found at either intergenic or exonic loci (**Fig 3B**). Genes bound by HNRNPM were surprisingly much longer on average (p<2.2e-16; **Fig 3C**), with median lengths of 3880bp, compared to the median lengths of 845bp in all Ensembl transcripts. HNRNPM-bound sites tended to be guanidine and uridine (GU) rich (**Fig 3D** and **3E**), and motif analyses of these sites revealed sequences consisting of single or double G residues interspersed with one to two U residues (**Fig 3E**).

**Figure 3:**
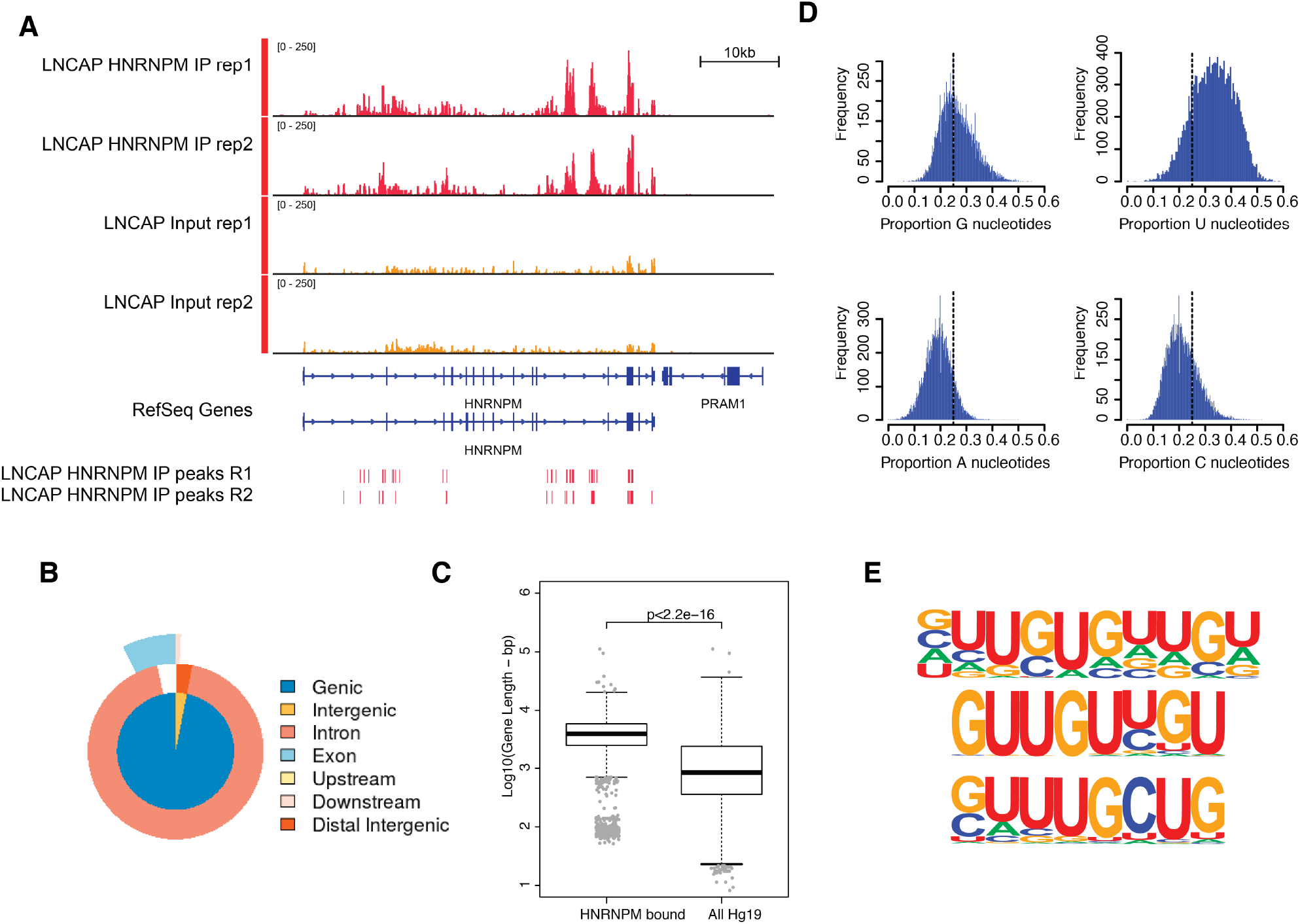
HNRNPM binds GU rich elements within long introns. **(A)** Normalized read density of HNRNPM eCLIP at the HNRNPM and PRAM1 genes. Input (yellow) and immunoprecipitation (red) tracks are shown. High confidence (p<0.05, Fold enrichment >2) peaks are shown in the lowest track. **(B)** Distribution of HNRNPM bound peaks across different genomic features **(C)** Distribution of gene lengths in all genes (blue) or HNRNPM bound genes (red) **(D)** Histogram plot of nucleotide frequencies within called, high confidence HNRNPM bound peaks. **(E)** Motif analysis of HNRNPM bound peaks. Shown are the top three (from top to bottom; p=1e-726, p=1e-625 and p=1e-597) motifs found within intragenic HNRNPM bound sites.

### Loss of HNRNPM results in minor changes in total mRNA levels

HNRNPM binding to pre-mRNA could potentially alter either transcript abundance or splicing. To better understand the impact of reduced HNRNPM binding in PCa cells, we further performed RNA sequencing on LNCAP PCa cells that were transduced either with the scrambled shRNA or HNRNPM-specific shRNAs (2B7 and 2B9) (**Fig S4A, Fig 4A, Table S4**). When compared to scrambled shRNA control treated cells, the overall transcript abundance shifts in both populations of HNRNPM shRNA-treated cells were positively correlated with each other (**Fig 4A**), suggesting that majority of the effects observed on the transcriptome were specific to a reduction in HNRNPM levels. Expression analyses revealed that: (1) only 0.58% (38 of 6510 mapped and bound genes) of HNRNPM-bound RNAs varied significantly (p<0.05, FDR <0.05, Fold Change >= 2 in both shRNA conditions) in expression levels during HNRNPM depletion in either knockdown condition (**Table S5**), and that (2) a minority (19.04%; 38 of 189 differentially and significantly expressed) of transcripts that were significantly altered in expression during HNRNPM knockdown were also bound by HNRNPM (**Fig 4A, Table S5**). These data suggested that the majority of transcript abundance changes occurring in these cells were not likely to be a direct consequence of HNRNPM depletion.

**Figure 4.**
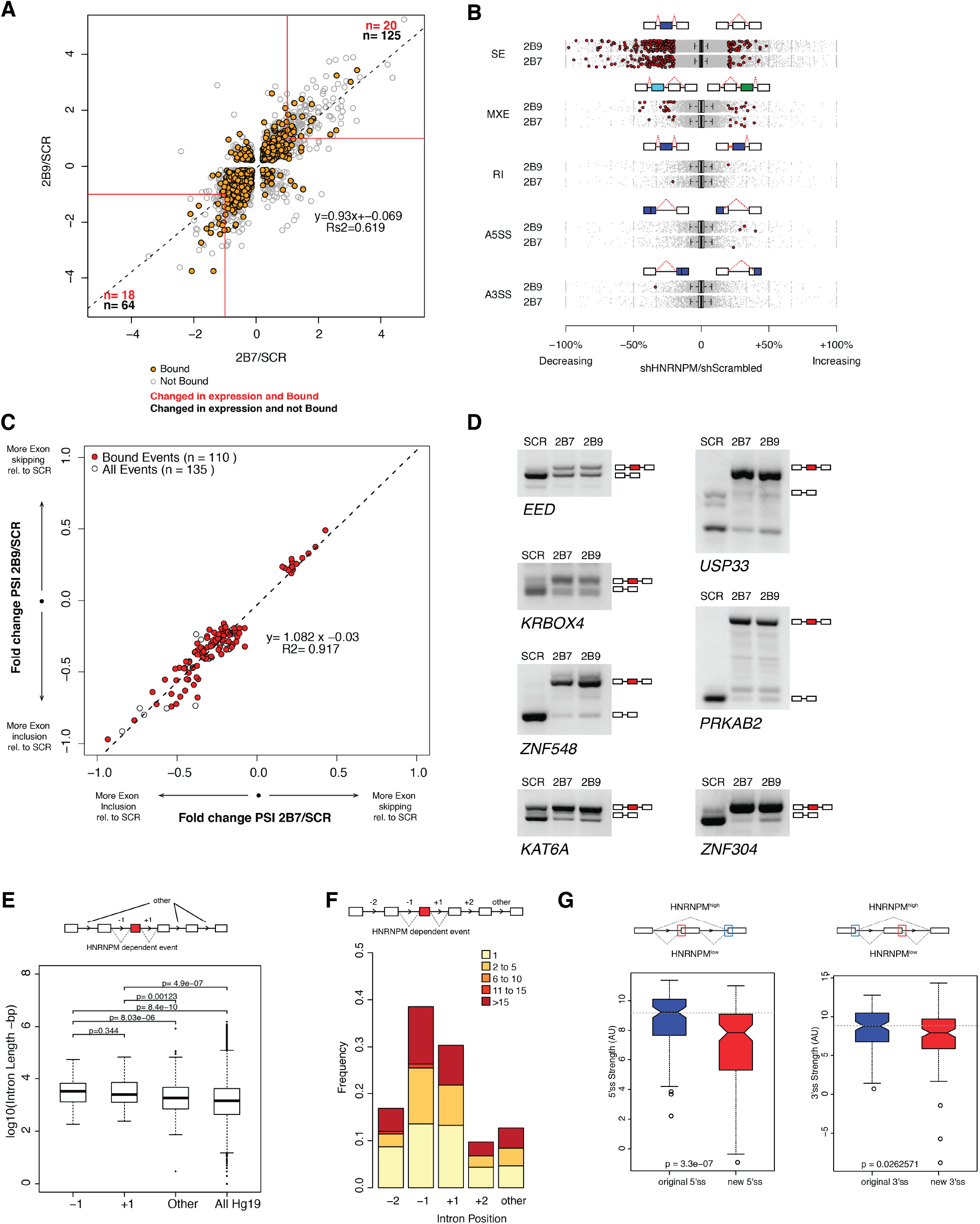
HNRNPM depletion results in increased exon inclusion. **(A)** Scatterplot of all significantly changed (p<0.05, FDR<0.05) genes (grey point) in HNRNPM shRNA treated cells, and their HNRNPM binding status. HNRNPM bound genes are highlighted in orange. Red boxes denote a 2 fold change cut off. The total numbers of differentially expressed genes (p<0.05, FDR<0.05, fold change >=2) are indicated in black, and the numbers of genes that are differentially expressed and bound by HNRNPM are indicated in red. **(B)** Plots showing all alternative splicing events occurring in 2B9 shRNA treated cells. All unique captured events are plotted in grey, while significantly changing events are shown in red. Change in percent spliced in (ΔPSI; HNRNPM shRNA (2B9) versus Scrambled shRNA) for the captured events is shown on the x axis. A similar plot for HNRNPM shRNA 2 (2B7) is shown in Figure S4b. RI: Retained Intron; A3SS: Alternative 3’ splice site usage; A5SS: Alternative 5’ splice site usage; MXE: Mutually exclusive exons; SE: Skipped Exon **(C)** Scatter plot of splicing events that are significantly changed (p<0.05, FDR<0.05, |ΔPSI| > 0.2) in both HNRNPM shRNA (2B7 and 2B9) conditions. Shown in red are events that occur in transcripts that are bound by HNRNPM. **(D)** RT-PCR validation of differentially spliced events that occur in HNRNPM depleted cells. **(E)** Distribution of unique intron lengths in the upstream (−1) or downstream (+1) introns flanking linear mis-splicing events compared to that of other introns in the same transcript or across the transcriptome. P-values were determined by the Wilcoxon test. **(F)** Distribution of the numbers of HNRNPM binding sites across the indicated introns in transcripts with linear mis-splicing events. **(G)** 5’ (right) and 3’ (left) splice site strength as compared to original, flanking splice sites in HNRNPM-dependent linear transcripts.

### Loss of HNRNPM results in increased exon inclusion

Alternative splicing of transcripts can give rise to events such as changes to alternative 3’ or 5’ splice site usage, cassette exon inclusion or exclusion or intron retention. To determine if some of these events were altered in HNRNPM depleted cells, we also performed splicing analyzes of our RNA sequencing dataset. These analyses revealed that the majority of significant splicing events affected by HNRNPM loss (p<0.05, FDR<0.05, dPSI>0.2) were the increased inclusion of cassette exons (**Fig 4B**). We observed few significant changes in alternative 5’ or 3’ splice site usage, in intron retention or in mutually exclusive exon usage in either shRNA conditions (**Fig 4B**).

As skipped exon (SE) events were the dominant alteration in HNRNPM depletion, we focused on this group of events for further analyses. We found a core group of 135 SE events that were present and significantly altered in both shRNA conditions. 110 (81.5%) of the core 135 identified events were found in genes that were bound by HNRNPM (**Fig 4C, Table S6**). Of these 110 events, 96 (87.3%) were events where the cassette exon displayed increased inclusion (**lower left quadrant**, **Fig. 4C**). We were able to independently validate several of the events (**Fig 4D**) via RT-PCR. Taken together, these data suggested that the majority of splicing changes occurring in HNRNPM-depleted cells were highly likely to be a direct consequence of reduced HNRNPM binding and that HNRNPM depletion tends to result in increased exon inclusion.

To better understand the specificity of HNRNPM towards its target introns, we examined the surrounding genomic and structural features of exons that were differentially included upon HNRNPM depletion. HNRNPM-silenced exons in linear transcripts were flanked by introns that were significantly longer (−1 flank median: 3327bp; +1 flank median: 2498bp; **Fig. 4E**) than other introns in the same transcripts (median:1826bp), or across the genome (median: 1434bp). HNRNPM binding sites were also preferentially found in these introns compared to other introns of the same transcript (**Fig. 4F**). Target exons appeared to bear weaker 5’ and 3’ splice sites compared to original flanking splice sites (**Fig 4G**). Overall, we can conclude that HNRNPM is enriched in the long proximal introns of mis-spliced exons.

### Loss of HNRNPM results in increased circular RNA formation

Aside from regulating alternative splicing events in linear mRNA, the splicing machinery is also known to catalyze the formation of exon-back splicing events in pre-mRNA to generate circular RNAs (CircRNAs). While the biological functions for most circRNAs remain unclear, their expression is highly regulated and specific to different tissue and cell types (Hansen et al., 2013; Memczak et al., 2013).

To determine if HNRNPM regulates circRNA biogenesis, we analyzed our RNA-sequencing dataset for backsplicing events. We identified a total of 1357 circularized transcripts in LNCAP cells amongst which 332 circRNAs that were significantly changed in expression. More circRNAs (230/340) were upregulated in HNRNPM deficient cells, suggesting that HNRNPM typically inhibits the formation of these RNAs (**Fig 5A and 5B, Table S7**). The vast majority of these circular RNAs were multi-exonic (312/340) (**Fig 5C**). We were able to validate several of these events in RNaseR treated RNA with RT-PCR using divergent PCR primers (**Fig 5D**). Similar to our observations for the linear events, we also observed that the majority (88.55%; 294/340) of transcripts that had changes in back-splicing were bound by HNRNPM (**Fig 5A**), suggesting that the differences in back-splicing frequency observed in HNRNPM-depleted cells is likely to be a direct consequence of reduced HNRNPM binding.

**Figure 5.**
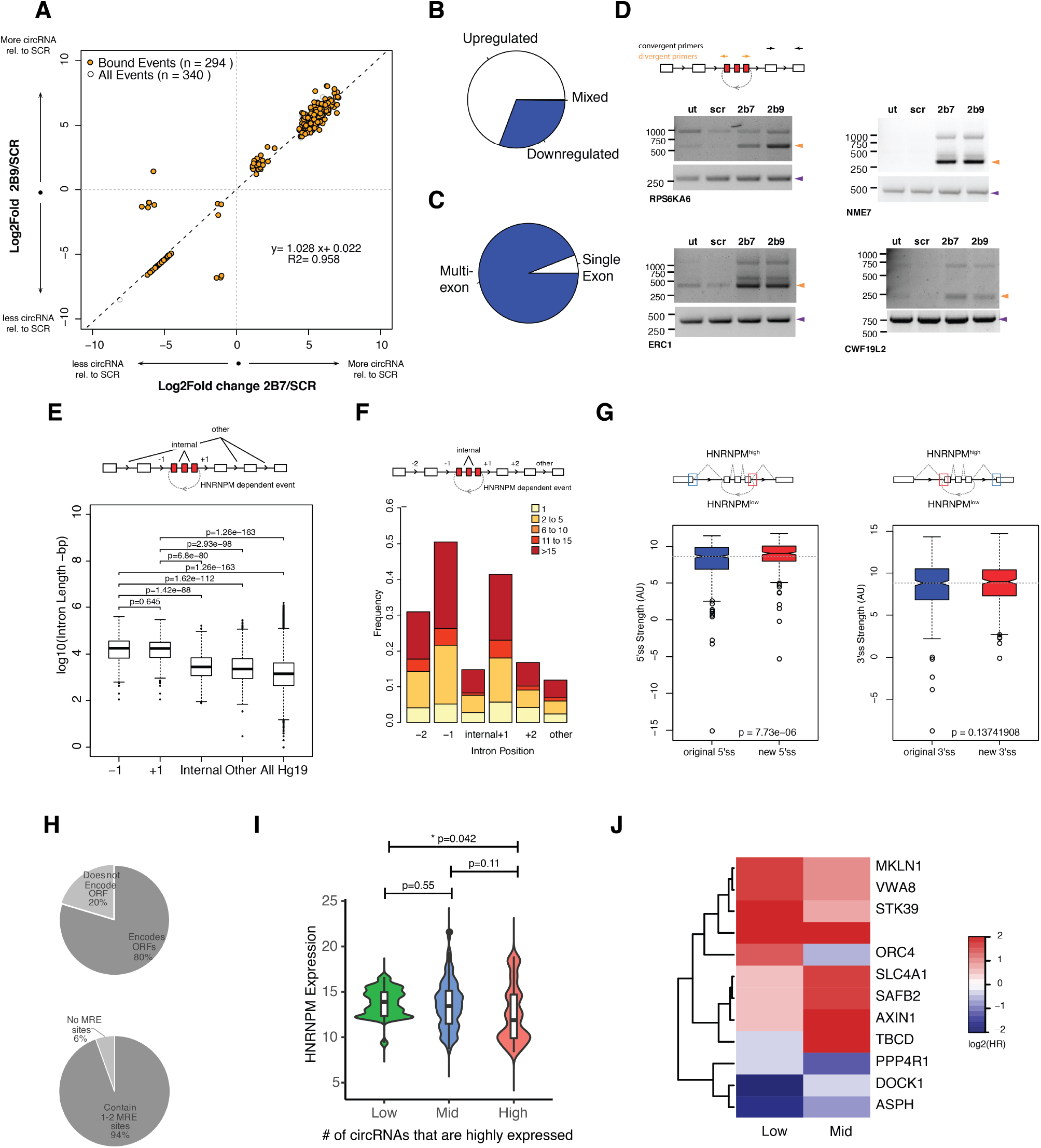
HNRNPM depletion results in increased circular RNA formation. **(A)** Scatterplot of CircRNA events that are significantly (p<0.05, FDR<0.05) changed in sh2B7 or sh2B9 treated cells compared to scrambled shRNA treated cells. All individual captured events are plotted in grey, genes highlighted in orange are bound by HNRNPM. **(B)** Distribution of upregulated and downregulated circular RNAs in HNRNPM deficient cells. **(D)** Distribution of multi-exonic or single exon circular RNAs regulated by HNRNPM. **(D)** Semi quantitative RT-PCR validation of circular RNA events occurring in sh2B7 or 2h2B9 treated cells. **(E)** Distribution of unique intron lengths in the upstream (−1) or downstream (+1) introns flanking circular mis-splicing events compared to that of other introns in the same transcript or across the transcriptome. P-values were determined by the Wilcoxon test. **(F)** Distribution of the numbers of HNRNPM binding sites across the indicated introns in transcripts with circular missplicing events. **(G)** 5’ (right) and 3’ (left) splice site strength as compared to original, flanking splice sites in HNRNPM-dependent circular transcripts. **(H)** Distribution of HNRNPM-dependent circular RNAs that encode ORFs or have miRNA-response elements. **(I)** HNRNPM expression levels in patients that express high, mid and low levels of target circRNAs **(J)** Heatmap showing the log2(Hazard Ratios) of patients expressing low or mid levels of the indicated circRNAs, when compared to patients expressing high levels of the same circRNA.

Similar to linear transcripts, introns flanking (−1 flank median: 17.5 kb; +1 flank median: 17.2 kb) HNRNPM-silenced circularizing exons were significantly longer than other introns (median:2320bp) in the same transcripts, or across the genome (**Fig 5E**). HNRNPM binding sites were also enriched in the proximal 5’ and 3’ introns flanking differentially circularized exons. Fewer HNRNPM binding sites were found in distal introns or introns within the circularized transcripts (**Fig 5F**). However, HNRNPM dependent back-spliced exons bore stronger 5’ and 3’ splice sites than that of the original flanking proximal exons (**Fig 5G**).

Finally, we tried to understand if HNRNPM dependent circRNAs would contribute to prostate cancer progression. Circular RNAs have been reported to regulate gene expression through microRNA (miRNA) response elements as miRNA sponge, through proteins encoded in their sequences. Circular RNAs regulated by HNRNPM had few miRNA binding sites, suggesting that they were not likely to perform as miRNA sponges because most of them contained only 1-2 common miRNA response elements (MREs). The majority of HNRNPM dependent circular RNAs (231/294) were however predicted to have coding potential (**Fig 5H, right panels**).

To then better understand the physiological relevance of these circRNAs, we took advantage of a recently published dataset (S. Chen et al., 2019) that analyzed circRNA expression in a cohort of 144 PCa patients. 248 of the 320 HNRNPM-regulated circRNAs were found expressed in patients. Expression of these circRNAs was also correlated with *HNRNPM* expression. Indeed, patients that had expressed higher levels of these circRNAs had significantly lower levels of *HNRNPM* (**Fig 5I**). Interestingly, low to mid expression of a cluster of HNRNPM-regulated circRNAs was significantly correlated with increased risk for patient biochemical relapse (BCR) (**Fig 5J and S4B**), and have a “tumor-suppressive” function. This was in-line with our observations that high *HNRNPM* levels, which are predicted to reduce the expression of these circRNAs, contributes to poorer prognosis in PCa.

### Structure-forming sequences are enriched in the flanking introns of mis-spliced events

To determine if HNRNPM binding was indeed sufficient to inhibit exon inclusion, we adopted the use of a bi-chromatic splicing reporter that expresses either GFP or dsRed fluorescent proteins, depending on the alternative splicing event that occurs (**Fig 6A**). HNRNPM binding sites of either the USP33 or APMAP transcripts were cloned into this reporter (**Fig 6B, Table S8**). As controls, we mutated predicted HNRNPM binding motifs (**Fig 3E**) found at each site (**Fig 6C**). Flow cytometry analysis of 293T cells transfected with either reporter showed that introduction of a HNRNPM binding site to the reporter resulted in increased RFP expression as compared to the empty vector control (**Fig 6D**). This suggests that, as predicted, the presence of a HNRNPM binding site in the reporter was sufficient to cause exon skipping. Conversely, mutation of the HNRNPM binding sites in the reporter resulted in increased GFP expression (**Fig 6D, right panels**). Taken together, these data add support for our observations that the presence of HNRNPM is indeed required to suppress exon inclusion in cells.

**Figure 6:**
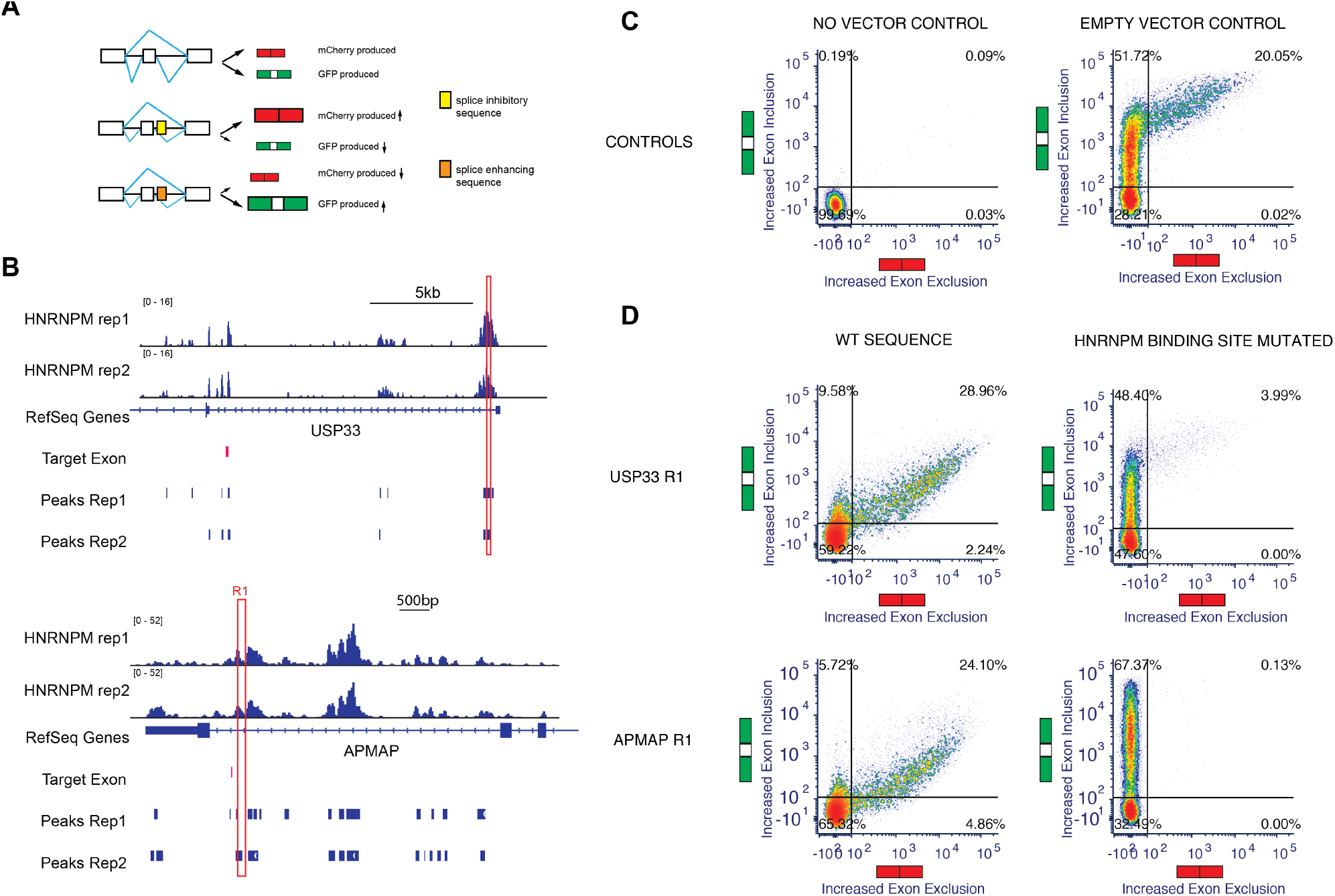
Genomic and structural features of HNRNPM-dependent silenced exons. **(A)** Schematic showing outcomes of minigene splicing assay **(B)** Plots showing HNRNPM bound regions of the USP33 and APMAP gene that were cloned and inserted into the minigene construct **(C)** Flow cytometry plots showing differential splicing patterns of cells that were transfected with the indicated constructs. Left: No plasmid control; Right: Unmodified vector control. Y-axis shows increasing levels in exon inclusion and x-axis shows increasing levels of exon skipping. **(D)** Flow cytometry plots showing differential splicing patterns of cells that were transfected with the indicated constructs. Left: Vectors with wildtype HNRNPM binding site, Right: Vectors with mutated HNRNPM binding site. Y-axis shows increasing levels in exon inclusion and x-axis shows increasing levels of exon skipping.

Overall, our observations suggest that HNRNPM preferentially binds to long introns in its target genes (**Fig 4F and 5E**), and that loss of binding in these introns results in accumulation of mis-splicing events. In long introns, RNA secondary and tertiary structures, stemming from competitive long and short range pairings between short inverted repeat (IR) sequences, has been shown to be partially required for appropriate splice site selection (Jeck et al., 2013; Rogic et al., 2008; Zhang et al., 2014). These pairings can be intra- or inter-intron, and may function to shorten effective branchpoint distance or mask cryptic splice sites. Such IR pairs tend to be derived from repetitive elements (short-interspersed nuclear elements; SINEs, long-interspersed nuclear elements; LINEs). We therefore examined if there was differential association of HNRNPM with these elements in mis-spliced transcripts but not in unaffected transcripts. Approximately 43.7% of HNRNPM bound peaks across the transcriptome were associated with at least one class of repetitive element (LINE, SINE, repetitive DNA, or long terminal repeats (LTR)) (**Fig S5A; All binding sites; bottom panel)**). In contrast, 54.1% and 50.5% of HNRNPM binding sites found within the proximal flanking introns of mis-spliced exons in linear and circular transcripts intersected with such elements (**Fig S5A; top left and middle panels**). Consistent with previous observations, L1 LINE elements represented the major class of repeats that HNRNPM peaks were associated with (**Fig S5B**; Kelley, Hendrickson, Tenen, and Rinn (2014)).

Recent studies have also suggested that HNRNP interactions with RNA tertiary structures such as RNA G quadruplexes (GQ) can impact splicing outcomes (Huang, Zhang, Harvey, Hu, & Cheng, 2017). Given that HNRNPM preferentially binds to GU-rich sequences, we also investigated if sequences found within flanking introns of mis-spliced events have the potential to form GQ structures. In linear transcripts, we observed an increased incidence of potential GQ forming sequences per kb of intron length in the intron upstream of the missplicing events (**Fig S5C; left panel**), compared to other introns in the same transcript. In contrast, potential GQ structures were enriched in both the upstream (−1) and downstream (+1) introns flanking mis-spliced circRNA events (**Fig S5C; right panel**). GQ, when complexed with hemin, has peroxidase activity that can be detected upon addition of substrate by a maximal absorbance at 420nm (Li et al., 2013). Using this assay, we were also able to confirm that several HNRNPM bound sequences had the capability to form tertiary GQ structures (**Fig S5D**). Examination of local secondary structure (+/- 1kb) by DMS-MaP sequencing around representative HNRNPM binding peaks in the flanking introns of five missplicing events in GMPR2, PRKAB2, USP33, ZNF548, ZNF304 also revealed the presence of many local structures at and around HNRNPM binding sites (**Fig S5E-G**). Formation of these structures however did not appear to be HNRNPM-dependent (**Fig S5G**).

### Mimicking the inclusion of HNRNPM-silenced poison exons by splice switching antisense oligonucleotides inhibits cell growth

Changes to alternative splicing can affect transcript fate and function(Ellis et al., 2012; Lareau & Brenner, 2015; Naro et al., 2017; t Hoen et al., 2011; Yang et al., 2016). This, in turn, leads to changes in both the cell’s transcriptome and proteome, manifesting as phenotypic transformations. Because the main impact of HNRNPM depletion were changes in alternative splicing, but not gene expression, we focused on the exon inclusion and circularization events that were changed upon HNRNPM knockdown. Gene ontology analysis of these genes suggested that most of these genes are involved in specific key cellular processes **(Fig 7A**), including histone lysine methylation and the regulation of microtubule polymerization.

**Figure 7:**
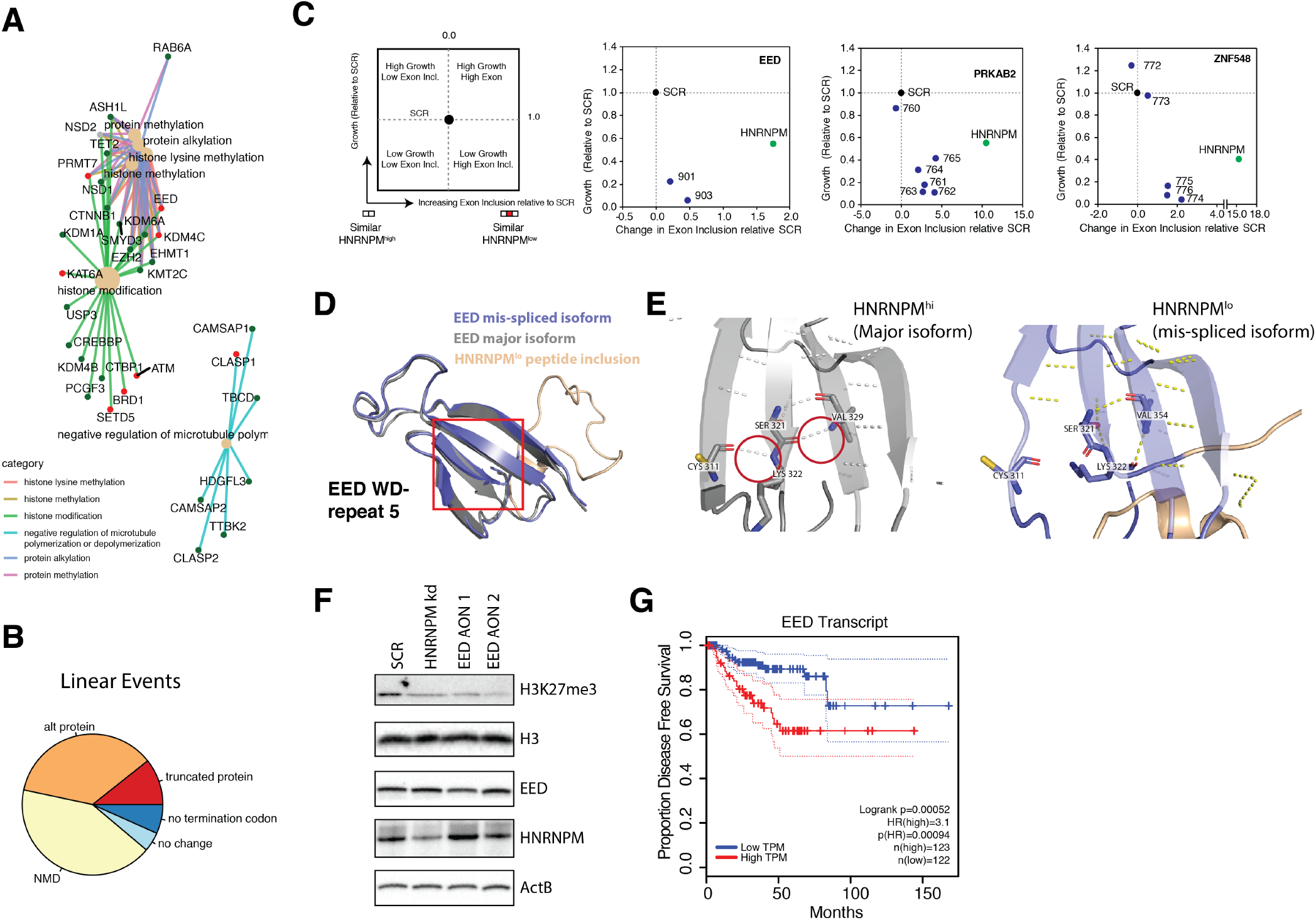
Mimicking HNRNPM-dependent linear splicing events in cells inhibits cell growth. **(A)** Gene-ontology analysis of transcripts that are either changed in expression or mis-spliced upon HNRNPM knockdown. **(B; leftmost Panels)** Schematic showing cell growth and exon inclusion outcomes in SSO treated cells as compared to scrambled SSO treated cells. (**B, right panels)** Distribution of predicted outcomes of HNRNPM-dependent linear splicing events occurring in coding domains of affected genes. **(C)** Scatter plots of overall cell growth yields (y-axis) in relation to overall exon inclusion levels (x-axis) upon treatment of the specified SSOs. Relative exon inclusion levels were measured and calculated based on target peak areas determined by FLA-PCR. Values are shown relative to cells treated with scrambled SSOs. Labels: SCR: Scrambled SSO; HNRNPM: HNRNPM-NMD inducing SSO; Numbered labels: Indices of gene-specific SSOs that induce inclusion of target HNRNPM^low^-dependent exons. A representative of two experiments is shown. **(D)** WD40 domain 5 of EED when the HNRNPM-regulated exon 10 is included. Structure of the HNRNPM regulated WD40 domain in wildtype EED (Grey), superimposed on the predicted structure of the HNRNPM-dependent EED isoform (blue) The new peptide generated by the splicing event is depicted in orange. **(E)** Close up view of hydrogen bonding interactions that stabilize the WD40 beta sheet in either wild-type or the new EED isoform. **(F)** Western blot of H3K27me3 (EED target) upon HNRNPM loss or EED event **(G)** Disease free survival curves of PCa patients when expression of EED is low (blue) or high (red)

The majority of HNRNPM-dependent linear splicing events were predicted to result in nonsense mediated decay (NMD) or alternative reading frames in their target protein (**Fig 7B**), suggesting that the functions or protein levels of many of these targets were likely to be altered in HNRNPM^low^ cells. To determine if the mis-splicing of some of these genes contributed to the reduction in cell growth, we designed splice switching antisense oligonucleotides (SSOs) to induce three HNRNPM-specific exon retention events in HNRNPM-sufficient LNCAP cells. Transfection of SSOs targeting splicing events occurring in *EED, PRKAB2*, and *ZNF548* recapitulated splicing events occurring in HNRNPM deficient cells (**Fig 7C**), and led to the loss of cell viability in target cells. As a control, we used an SSO against HNRNPM that is predicted to result in NMD of the HNRNPM transcript. Our results suggest that the reduction in cell growth in HNRNPM deficient cells is likely a result of perturbing multiple genes across different pathways.

### EED isoform 2 is a novel splicing target of HNRNPM

While HNRNPM loss resulted in NMD events, a significant portion of them also resulted in alternative proteins (**Fig 7B**). One such event occurred in the *EED* transcript, which our SSO data suggested was growth inhibitory. We were interested in understanding why this was so. Loss of HNRNPM expression resulted in increased inclusion of *EED* exon 10, resulting in a 25-amino acid added in-frame to the EED protein. To determine how this may affect EED function, we analyzed and modelled the structure of EED protein with or without the peptide extension. Wildtype EED contains seven copies of the WD-repeat motif, and folds into a seven-bladed beta-propeller structure. We found that addition of the peptide sequence into the structure results in disruption of hydrogen bonding within the beta sheet of WD repeat 5 (**Fig 7D, 7E and S6A**). This is likely to disrupt the stability of the WD repeat, and affect EED function. In support of this, we find that induction of this specific isoform of EED in LNCAP cells is correlated with reduced H3K27me3 activity (**Fig 7F**). These results suggest that loss of EED function is likely to contribute to the reduced proliferation we observed upon HNRNPM. Indeed, analyses of the TCGA data sets suggest that patients with reduced EED expression have improved disease-free survival, compared to patients with high EED expression (**Fig 7G**).

## DISCUSSION

Recent evidence suggests that cancer cells may be especially vulnerable to disruptions in their splicing machinery than normal, untransformed cells (Braun et al., 2017; Hsu et al., 2015; Koh et al., 2015). This is thought to be a consequence of increased proliferative rates and transcriptional outputs, resulting in “addiction” to a highly functioning spliceosome. Despite this, splice signatures amongst different cancers can vary (Sebestyen, Zawisza, & Eyras, 2015), suggesting that unique or sets of splicing factors may regulate different splicing programs across different cancer types. For instance, ESRP1 has been suggested to drive both oncogenic (Fagoonee et al., 2017) and tumor suppressive (Shapiro et al., 2011; Ueda et al., 2014) programs in the context of different cancers. Identification of key factors that underpin a cancer-type specific splicing program is, therefore, valuable in the development of downstream therapeutics.

To this end, we have, through an unbiased and systematic screen of splicing factors, identified HNRNPM as a potential vulnerability in prostate cancer (Fig 1). HNRNPM depleted LNCAP prostate cancer cells fail to expand both *in vitro* and *in vivo* (Fig 2). Our data suggest that the arrest in cell proliferation in HNRNPM-depleted cells is likely a consequence of missplicing, as opposed to an alteration of transcript abundance. Indeed, using antisense oligonucleotides to mimic several of these splicing events in cells that expressed HNRNPM was sufficient to inhibit cell proliferation and growth (Fig 7C). In support of this, we showed that an exon inclusion event in the EED transcript results in destabilization of one of its WD-repeats, and is likely to contribute to reduced PRC2 complex activity in PCa cells (Fig 7D-G). We have also showed that the majority of circRNAs regulated by HNRNPM are expressed in PCa patients, and that their expression levels are anti-correlated with that of HNRNPM (Fig 5I). Reduced expression of these circRNAs resulted in poorer disease-free survivals in these patients (Fig 5J). Taken together, HNRNPM likely regulates PCa cell proliferation by suppressing exon inclusion and circularization of multiple transcripts across many key homeostatic pathways in cells.

Interestingly, the phenotypic manifestation of HNRNPM loss in prostate cancer cells is different from what was previously described. Here we describe a role for HNRNPM in maintaining cell fitness but did not observe changes in cell identity programs in PCa cells. In breast cancer cells, where loss of HNRNPM inhibits epithelial-mesenchymal transition (EMT), but not cell growth(Xu et al., 2014). Together, these data are consistent with a contextdependent function for HNRNPM in cancer, which might depend on its downstream targets. Interestingly, the herein identified downstream target EED controls cell proliferation and EMT in context-dependent manner(Serresi et al., 2016; Shi et al., 2013).

Splicing defects occurring during HNRNPM deficiency primarily involve increased exon inclusion and exon circularization, indicating that HNRNPM normally functions to suppress splice site usage in PCa. This is consistent with previous observations on splicing during HNRNPM loss in other cell types (Damianov et al., 2016; Huelga et al., 2012; Xu et al., 2014). Our data indicate that mis-spliced linear-splicing and circularization events occurring during HNRNPM deficiency are located in exons that are flanked by HNRNPM-bound introns. HNRNPM occupancy is much lower in other introns in the same transcripts, suggesting that its recruitment is important for ensuring appropriate splice site selection.

Members of the hnRNP family of proteins have been shown to be important in ensuring correct splice site selection in target genes. These proteins tend to interact with cis-regulatory splicing silencer sequences in pre-mRNA, and several hnRNPs have also been identified as repressors of splicing (Matlin, Clark, & Smith, 2005). hnRNP-mediated splicing repression has been attributed to several modes of action, including antagonizing core spliceosomal components and/or splicing enhancer proteins from binding to pre-mRNA (Mayeda, Helfman, & Krainer, 1993; Mayeda & Krainer, 1992; Zhu, Mayeda, & Krainer, 2001), as well as alteration of long-range secondary structure in pre-mRNA to inhibit/promote alternate splice site usage (Blanchette & Chabot, 1999; Martinez-Contreras et al., 2006; Solnick & Lee, 1987). These modes of action are not mutually exclusive, and some hnRNPs (e.g. hnRNPA1) have been shown to rely on both, depending their binding context (Blanchette & Chabot, 1999; Martinez-Contreras et al., 2006; Mayeda et al., 1993; Mayeda & Krainer, 1992).

Our analyses suggest that HNRNPM specifically prevents mis-splicing in the context of long genes and is especially enriched in long introns across the genome. Increased intron length has been shown to adversely affect correct splice site selection. For one, cryptic splice sites that compete with the original 5’ and 3’ splice sites are more likely to be found within a longer intron. Secondly, increased intron length extends the delay between RNA polymerase II-dependent synthesis of the 5’ and its associated 3’ splice sites, resulting in a higher probability of RNA secondary and/or tertiary interactions forming to bridge a 5’ss and an incorrect proximal cryptic 3’ss.

Formation of RNA secondary and tertiary structures has been shown to be partially required for correct splice site selection in long introns (Rogic et al., 2008; Zhang et al., 2014). In vertebrates, large introns bear relatively high densities of repetitive regions that are predicted to form both intra- and inter-intron complementary base pairings (Shepard, McCreary, & Fedorov, 2009). Such pairings may be long range ( >1.4 kb apart; Aktas et al. (2017)) and are typically formed between long or short interspersed repetitive elements (LINEs and SINEs). Competition between such pairs of complementary sequences has been shown to significantly alter splice site selection in both linear and circular transcripts (Zhang et al., 2014). It may be that HNRNPM is required to regulate the formation or interactions between these structures at such transcripts, in order to maintain correct splice site selection. In line with this, HNRNPM was previously shown to interact with several ATP-dependent RNA helicases (Brannan et al., 2016), pointing to a possible association with RNA secondary/tertiary structure. We also observe that HNRNPM is more enriched at repetitive elements in mis-spliced transcripts than other bound transcripts. Long flanking introns that flank mis-splicing events in HNRNPM deficiency also have a higher tendency to form G-quadruplex structures. Interestingly, we do not observe local (<1kb flanking HNRNPM bound sites) changes in secondary structure immediately surrounding HNRNPM bound sites. It could be that HNRNPM is more important for mediating cross-intron interactions or required for regulating RNA tertiary structures.

Finally, it has been previously reported that long, alternatively spliced genes, bearing elevated intron length to exon length ratios, are highly enriched in pathways associated with cancer and other multi-genic diseases (Sahakyan & Balasubramanian, 2016). Understanding how this class of genes is regulated may thus be critical for developing therapeutics against these diseases. Because of their lengths, expression of this class of genes can be rate-limiting for cell survival and/or function. Inhibition of transcription elongation by blocking topoisomerase activity is sufficient to impair their expression (King et al., 2013). In a similar vein, we have shown that expression of these genes may also be especially vulnerable to depletion of splicing factors such as HNRNPM. One prediction that arises from this observation is that expression of these genes may be modulated through synergistic interactions between inhibitors targeting transcription elongation and splicing factors. Indeed, there is some evidence in the literature pointing towards this. Mutations in the splicing factors SRSF2 and U2AF1 have been shown to impair transcription pause-release and drive R loop formation (L. Chen et al., 2018), suggesting that these two pathways are intimately linked. Identification of other factors similar to HNRNPM that may similarly affect homeostasis of these genes can thus be important for cancer therapy.

## ACKNOWLEDGEMENTS

We thank A Jeyasekharan, M Hoppe for sharing protocols and helpful discussions. We are grateful to the staff at the A*STAR Biological Resource Center for support for animal work, the GIS Genome sequencing team for help with RNA sequencing, and the entire E.G laboratory for critical discussion. This work was supported by the Institute of Molecular and Cell Biology, Agency for Science Technology and Research, Singapore. EG acknowledges support from NMRC/OFIRG/0032/2017, NRF-CRP17-2017-06 and MSSM seed fund. Research reported in this publication was additionally supported by the Tisch Cancer Institute through the National Cancer Institute Cancer Center Support Grant (P30 CA196521). G.G. lab is supported by the MDC, the Helmholtz Association and the ERC.

## AUTHOR CONTRIBUTIONS

Overall design of the project: JH and EG. Acquisition of experimental data: JH, MS, DI, FG, TT, HW, AHH, SZ. Computational and genomics approaches: DL, DI, JZ, LC, OA, THMC, VR, BDG, SO, DKBW, SC, MA, HHH. Generation of reagents and scientific inputs: MS, GG, SC, MA, HHH, IM, DKBW, EG. JH and EG wrote the manuscript with comments from all authors.

## COMPETING FINANCIAL INTERESTS

The authors declare no competing interests

## DATA AVAILABILITY

All data needed to evaluate the conclusions in this study are present in the paper and/or its Supplementary Materials. eCLIP and RNA-Sequencing data supporting the findings of this study have been deposited into the National Center for Biotechnology Information (NCBI) Gene Expression Omnibus under accessions GSE113786.

## MATERIALS AND METHODS

### Cell Culture and treatment

LNCAP (ATCC) and PC3 (ATCC) cells were cultured and maintained in RPMI media supplemented with 10% fetal bovine serum (FBS; Lonza) 1mM sodium pyruvate (Gibco, 11360070), non-essential amino acids (Gibco; 11140050) and 50U/ml Penicillin-Streptomycin (Gibco, 15140163). Primary prostate epithelial cells (PrEC; Lonza, CC-2555) were cultured in PrEGM Prostate Epithelial Cell Growth Medium (Lonza, CC-3165) supplemented with BPE, Hydrocortisone, hEGF, Epinephrine, Transferrin, Insulin, Retinoic Acid, Triiodothyronine and GA-1000 as per the manufacturer’s recommendation (Lonza, CC-4177). HEK293T (ATCC) cells were maintained in DMEM media supplemented with 10% FBS, 1mM sodium pyruvate, non-essential amino acids, and 50U/ml Penicillin-streptomycin.

### Lentivirus production and infection

For single lentivirus generation, HEK293T cells were seeded to 50% confluency one day prior to transfection with the PLKO.1 vector of interest, packaging (d8.9) and envelope plasmids (VSVg) in the ratio of 5:5:1, using Lipofectamine 2000 reagent (Invitrogen). Fresh media was added to the transfected cells after an 18h incubation. Subsequently, cell culture supernatant containing lentiviral particles was collected at both 24h and 48h post media-change, filtered with a 0.22 μm filter, and concentrated by ultracentrifugation at 4°C, 23000 RPM for 2 hours. For pooled lentivirus generation, PLKO.1 vectors of interest were pooled prior to transfection in HEK293T cells with packaging and envelope plasmids. Ratios of vectors used were maintained as stated in the procedure used for single lentivirus production.

### shRNA pooled screen

The screens were conducted as previously described (Gargiulo, Serresi, Cesaroni, Hulsman, & van Lohuizen, 2014), with modifications. 5E6 LNCAP cells were seeded one day prior to infection in 150mm dishes, and infected with lentivirus pools at a multiplicity of infection of 1, in the presence of 8ug/ml polybrene. After a 48hour infection, cells were selected in 1ug/ml puromycin for 4-5 days. Upon completion of selection, cells were trypsinized and counted. 1.5E6 cells were harvested and saved as the input control for sequencing. The remaining cells were used either for in vitro passaging or in vivo tumorigenesis experiments. For the *in vitro* passaging experiments, 1.5E6 cells were plated in a 150mm dish and cells were harvested every 3-4 population doublings, with 1.5E6 cells harvested and plated at each passage. Harvested cell pellets were flash frozen and stored at −80C. For each injected tumor in the in vivo tumorigenesis experiments, 1.5E6 cells were re-suspended in 100ul of cell culture media mixed with Matrigel in a 1:1 ratio. Cells were then injected into the flanks of 6-8 week old, male CB17-SCID mice (In Vivos) and allowed to form tumors over time. Mice were monitored for tumor growth every 2-3 days. Tumors were harvested when they attained a size of 400mm^3^ and above. Harvested tumors were minced, trypsinized and ran through a 40um mesh to form single cell suspensions. Cell pellets were then flash frozen and stored at −80C. Genomic DNA from either the in vitro passaged or the in vivo tumor cell pellets was then harvested with the QIAGEN DNeasy kit in accordance to the manufacturer’s protocol. For lentivirus input libraries, RNA was isolated from ~3.5E6 infectious units of lentivirus using a combination of Trizol reagent (Invitrogen) and the Ambion Purelink RNA isolation kit. RNA was converted into cDNA using the Invitrogen Maxima cDNA synthesis reagent. shRNAs were amplified from genomic DNA or cDNA using primers spanning the common flanks of the shRNA expression cassette. These primers included adaptors for Illumina Hiseq sequencing. After PCR amplification, the constituent hairpins in each sample were identified through high-throughput sequencing, and the fold change in hairpin representation was determined by comparing the normalized reads of each sample to the relevant input (P1), plasmid and viral samples.

### Mice

Mice were housed in compliance with the Institutional Animal Care and Use Committee (IACUC) guidelines. All procedures involving the use of mice were approved by the local Institutional Animal Care and Use Committee (IACUC) and were in agreement with ASTAR ACUC standards.

### Short term siRNA screen

1E4 PrEC cells per well in a 384 well plate were reverse transfected with siRNA using Lipofectamine RNAimax reagent (Invitrogen), to a final concentration of 25nM. siRNAs against PLK1 were used as a positive control, whereas control non-targeting siRNAs were used as negative controls. 96 hours post transfection, total cell yields were measured using the CellTiter 96^®^ AQueous One Solution Cell Proliferation Assay (MTS), as per the manufacturer’s instructions. Absorbance readings (490nm) were taken at 2hours post incubation with the MTS solution. The relative impact of knockdown on cell growth was then determined by comparing absorbance readings from each sample well to that of the PLK1 or control non-targeting siRNA treated wells.

### RNA extraction and RT-PCR

For RNA was isolated using the Purelink RNA Mini Kit (Thermo Fisher Scientific; 12183018A)in conjunction with Trizol Reagent (Invitrogen; 15596018) following the manufacturer’s protocol. Following reverse transcription to cDNA with the Maxima First Strand cDNA synthesis kit (Thermo Fisher Scientific; K1641), 30ng of cDNA was used for each RT-PCR reaction.

### Circular RNA validation

RNA was purified as described above. After RNA isolation, 1ug of RNA was incubated with 5U of RNaseR (Epicentre) for 45 min at 37C in a 14ul reaction volume to digest all linear RNA. After digestion, RnaseR treated RNA was reverse transcribed to cDNA using the Maxima cDNA synthesis kit, as per the manufacturer’s recommendation. Semi quantitative RT PCR was then performed on the cDNA using divergent primers to determine levels of circRNA across samples. CircRNA exon-exon joints were further confirmed by excising PCR amplified bands in the gel and performing sanger sequencing.

### Western blot analyses

Cells were lysed in RIPA buffer (150mM sodium chloride, 1% NP-40, 0.5% sodium deoxycholate, 0.1% SDS, 50mM Tris pH 8.0) for 30min on ice, before being subjected to sonication (10sec, Setting 4, Microson XL2000). 10-30ug of protein lysates were ran per lane in 10% SDS-PAGE gels for 1h 15min at 125V. Proteins were transferred onto 0.22um nitrocellulose membranes by wet transfer (Tris-Glycine buffer, 20% Methanol) at 30mA for 1-2hours on ice. Membranes were blocked in 3% milk, before being incubated in the indicated antibodies overnight. Primary antibody bound membranes were then washed for 5 min three times in PBS+0.1% Tween-20, before being probed with HRP conjugated secondary antibodies for 45 min at room temperature. After three washes in PBS+0.1% Tween-20, membranes were probed with ECL, and imaged using film.

### eCLIP

eCLIP experiments were performed as described in Van Nostrand et al. (2016), but with the following modifications: ~2.5E7 LNCAP cells were grown on 150mm dishes were UV crosslinked (150mJ), using the UV Stratalinker 2400. Cell pellets were flash frozen in liquid nitrogen and stored at −80C for > 1 day to facilitate lysis. Lysis was performed in 1ml ice cold iCLIP lysis buffer (50mM Tris-HCL pH7.5, 100mM NaCl, 1% Igepal CA630, 0.1% SDS, 0.5% Sodium deoxycholate and 1.1% Murine Rnase Inhibitor (M0214L, NEB). Cells were allowed to lyse for 15 min on ice, before a short sonication (3 min; 30 sec on / 30 sec off; low setting) in the Diagenode Bioruptor. Sonicated cell lysates were then treated with 40U of RnaseI (Ambion) and 4U of Turbo Dnase (Ambion) for 5 min at 37C. RNase activity was inhibited with the addition of 1% SuperaseIn reagent, before being clarified with a 15000g centrifugation for 15 min at 4C. Clarified lysates were then subjected to an overnight IP at 4C, using 10ug of HNRNPM antibody bound to Protein G dynabeads. 10% input samples were set aside prior to the addition of antibody-bead complexes.

After the overnight IP, immunoprecipitated Protein-RNA complexes were treated with FastAP and PNK treated in the presence of SuperaseIn Rnase Inhibitors. This was followed by the ligation of a 3’RNA linker. The protein-RNA complexes were ran on a 1.5mm 4-12% NuPage Bis-Tris gel at 150V for 75min before being transferred on a 0.22um nitrocellulose membrane (30mA, 2hours). After the transfer, membranes were washed once in PBS. The regions corresponding to approximately 10KDa below and 75KDa above HNRNPM bound complexes were then excised. The region that was excised in this step was further confirmed to contain HNRNPM by running a separate western blot using aliquots of the same lysates. Excised membrane pieces were subjected to proteinase K digestion and RNA isolation in acid phenol/chloroform. The Zymo RNA Clean and Concentrator kit was used to purify the RNA containing aqueous layer from the phenol-chloroform extraction.

After purification, extracted RNA was reverse transcribed, and a 5’ linker was ligated in an overnight DNA ligase reaction. Linker ligated cDNA was then subjected to clean up, and amplified using PCR. The final library was purified using 1.8X Ampure XP beads. Adapter dimers were removed using a second Ampure XP bead selection step with 1.4X bead to eluate ratio. Library sizing was checked using the DNA high sensitivity Bioanalyzer chip, and libraries ranged from 150-300bp in size. Final libraries were pooled and sequenced using the Illumina HiSeq.

### SSO transfections

For proliferation assays, 8E3 LNCAP cells per well in a 96 well plate were reverse transfected with SSOs using the Lipofectamine RNAimax reagent to a final concentration of 100nM. Control, non-targeting SSOs were used as controls. Total cell yields were measured using the CellTiter 96^®^ AQueous One Solution Cell Proliferation Assay (MTS) at 48 hours post transfection, as per the manufacturer’s instructions.

Since the majority of the splicing events we were interested in were deleterious to cells, as compared to the non-targeting controls, reduced concentrations of SSOs were used in order to lower overall transfection efficiency and recover sufficient RNA and protein for downstream analyses. 3E5 LNCAP cells were reverse transfected with SSOs to a final concentration of 25nM, and harvested at 48hours post transfection. For protein assays, cell pellets were lysed in RIPA buffer containing protease inhibitors, before being briefly sonicated. After sonication, lysates were centrifuged (14000rpm, 4’C, 10 min) to remove cellular debris, and the resulting supernatant was quantified and used in western blot assays. RNA isolation was performed as described above.

### Bioinformatics analysis

RNA sequencing reads were mapped to the hg19 genome using STAR v2.4.2a (Dobin et al., 2013). Differential expression analysis was done using the Cufflinks suite v2.2.1 (Trapnell et al., 2012), and alternative splicing analysis was done using rMATS 3.0.9 (Shen et al., 2014), both using the Ensembl release 72 human annotation. Downstream splicing predictions was carried out using SPLINTER (Low, 2017).

e-CLIP-seq analysis was performed according to the protocol outlined in Van Nostrand et al. (2016)with several modifications outlined here. Briefly, pseudo paired-end reads were first created from single-end reads using bbmap in the BBTools suite (https://jgi.doe.gov/data-and-tools/bbtools/). The first 6bp of reads were also removed in the adapter-trimming step to improve alignment quality. Subsequent downstream analyses post-protocol were carried out using in-house R scripts.

### CircRNA expression and HNRNPM expression in PCa Patients

Patient clinical attributes and circRNA expression levels were obtained from Ref (S. Chen et al., 2019). A circRNA was considered as highly expressed in a patient if it was expressed at levels that were at the 75^th^ percentile of that of the cohort. To calculate HNRNPM expression levels in different groups of patients, the number of highly expressed circular RNAs were first counted per patient. Patients were stratified by those that had most numbers of high expressed circRNAs (>75^th^ percentile of cohort; Group=High), least (≤25^th^ percentile of cohort; Group=Low) or Mid-range (25^th^-75^th^ percentile of cohort; Group=Mid). HNRNPM levels for each group was then plotted.

### CircRNA expression and BCR Survival in PCa Patients

Patient clinical attributes and circRNA expression levels were obtained from Ref (S. Chen et al., 2019). For each circRNA, [atients were stratified by those that had high expression circRNAs (>75^th^ percentile of cohort; Group=High), low expression (≤25^th^ percentile of cohort; Group=Low) or Mid-expression (25^th^-75^th^ percentile of cohort; Group=Mid). Biochemical Relapse survival curves and hazard ratios were then calculated using the ‘Survival’ package in R.

### Fragment Length Analysis PCR (FLAPCR)

30ng of cDNA was amplified using 2X DreamTaq Mastermix (ThermoFisher, EP1701) and primers specific to the splicing target of interest. Forward primers were FAM-labelled. PCR products were amplified using the following program: 94°C for 5 minutes, then 30 3-steps cycles (94°C for 30”, 56°C for 45” and 72°C for 45”) and subsequent 8 3-steps cycles (94°C for 30”, 52°C for 45” and 72°C for 45”) with a final cycle at 72°C for 3 minutes. 1ul of the PCR product was added to 9.1 ul of HiDi + Liz markers premix (GeneScan 500/1200 LIZ dye Size Standard, ThermoFisher #4322682/4379950, optimized to give 1K RFU peak height) to each sample. The mix was denatured at 96°C for 3 minutes, cooled at 4°C for 3 mins and run in the ThermoFisher 3730XL DNA Analyzer. GeneMapper 5 (ThermoFisher) software was used to analyze the peaks.

#### Splicing Reporter Assays

Bichromatic fluorescent splicing reporter plasmid (RG6) were previously described in Ref (Orengo, Bundman, & Cooper, 2006) and obtained from Addgene (Plasmid #80167). Briefly, RG6 was digested with EcoRI and XhoI. Inserts corresponding to HNRNPM binding regions for APMAP1 (chr20:24945080-24945163; Hg19) and USP33 (chr1:78224956-78225074; Hg19) were cloned in. Sequences are shown in Table Sx. Where necessary, HNRNPM binding sites were mutated using QuikChange XLII Mutagenesis Kit (Stratagene). For flow cytometry assays, 2.5ug of each plasmid was transfected into 5E5 HEK293T cells. 2 days post transfection, cells were trypsinized and strained through a 40uM filter. Flow cytometry experiments were performed on the BD LSRII. Only singlet and live cells (as determined by DAPI dye exclusion) were analyzed. Analysis was performed with FCS Express 6.0.

#### *In vivo* DMS probing

LNCaP cells were trypsinized for 1 minute at 37°C, trypsin was neutralized by addition of fresh serum-containing medium, and harvested by centrifugation at 4°C for 5 minutes. After aspiring medium, cells were washed once in PBS, and resuspended in DMS Probing Buffer [10 mM HEPES-KOH pH 7.9; 140 mM NaCl; 3 mM KCl]. DMS was pre-diluted 1:6 in ethanol, then added to the cell suspension to a final concentration of 100 mM. Cells were incubated at 25°C with moderate shaking for 2 minutes. Reaction was quenched by placing cells in ice, and immediately adding DTT to a final concentration of 0.7 M. Cells were briefly collected by centrifugation for 1 minute at 9,600*g* (4°C). Supernatant was aspired, and cells were washed once with a solution of 0.7 M DTT in PBS.

#### Isolation of nascent RNA

The pellet from a 75%-confluence 100mm plate was resuspended in 150 μl of Cytoplasm Buffer [10mM Tris HCl pH 7; 150 mM NaCl] by pipetting using a cut P1000-tip. 50 μl of Cytoplasm Lysis Buffer [0.6% NP-40; 10mM Tris HCl pH 7; 150 mM NaCl] were slowly added dropwise, after which sample was incubated on ice for 2 minutes. The lysate was then layered on top of a 500 μl cushion of Sucrose Buffer [10mM Tris HCl pH 7; 150mM NaCl; 25% Sucrose]. Sample was then centrifuged at 16,000*g* for 10 minutes (4°C). The supernatant (representing the cytosolic fraction) was discarded, and the nuclei were washed in 800 μl of Nuclei Wash Buffer [0.1% Triton X-100; 1mM EDTA in 1X PBS], then collected by centrifugation at 1,150*g* for 1 minute (4°C). After removing the supernatant, nuclei were resuspended in 200 μl of Glycerol Buffer [20mM Tris HCl pH 8; 75mM NaCl; 0.5mM EDTA; 50% Glycerol; 10mM DTT] and resuspended by pipetting, then transfered to a clean tube. 200 μl of Nuclei Lysis Buffer [1% NP-40; 20mM HEPES pH 7.5; 300mM NaCl; 2M UREA; 0.2mM EDTA; 10mM DTT] were then added dropwise, while gently vortexing the sample. The sample was then incubated on ice for 20 minutes, then centrifuged at 18,500*g* for 10 minutes (4°C). The supernatant (representing the nucleoplasm) was discarded, and the chromatin fraction was washed once with Nuclei Lysis Buffer, without letting the pellet detach from the bottom of the tube. After discarding the supernatant, 1 mL of ice-cold TRIzol was added directly to the chromatin fraction, and the sample was incubated at 56°C for 5 minutes to solubilize chromatin. RNA extraction was performed by adding 200 μl of Chloroform, vigorously mixing for 15 seconds, then centrifuging at 12,500*g* for 15 minutes (4°C). After centrifugation, the upper aqueous layer was transferred to a new tube, and mixed with 3 volumes of ethanol by vigorously vortexing for 15 seconds. The sample was then loaded on a Zymo RNA Clean & Concentrator-5 column, and purified following manufacturer instructions.

#### RNA G-quadruplex Hemin Assays

Hemin assays for G quadruplex formation were conducted as previously described (Zheng et al., 2015). Briefly, RNA oligos corresponding to wildtype or mutated sequences of HNRNPM binding sites found in the introns of EED, USP33 and PRKAB2 transcripts were ordered from IDT DNA. The sequences of these oligos are found in **Table S9.** RNA oligos were first suspended to 10uM in folding buffer (20 mM Tris pH 7.6, 100 mM KCl, 1 mM EDTA). They were then heated to 98°C in a water bath and allowed to come to room temperature through passive cooling. Folded RNAs were incubated with 12uM of Hemin (Sigma H-5533, in DMSO) in 1X Buffer 2 (New England Biolabs) at 37°C for 1 hr. Substrate solution (2 mM ABTS, Sigma A9941; 2 mM hydrogen peroxide, Sigma H1009; 25 mM HEPES pH7.4, 0.2 M NaCl, 10 mM KCl, 0.05% Triton X-100, 1% DMSO) was then added to each mixture. After 15min, absorbance of each sample was measured from 400nm to 500nm using a SpectraMax Plus384 Microplate Reader. Samples containing RNA G-quadruplexes display the characteristic absorbance peak at 420nm.

#### Targeted DMS-MaPseq

DMS-MaPseq was performed with minor changes to the original protocol (Zubradt et al., 2017). Oligonucleotides were designed to amplify regions not exceeding a length of 600 nt. Reverse transcription was carried out in a final volume of 20 μl. 1 μg of nascent RNA was initially mixed with 2 μl of 10 mM dNTPs and 1 μl of 10 μM gene-specific primers, then denatured at 70°C for 5 minutes, after which the sample was allowed to slowly cool to room temperature by incubation for 5 minutes on the bench. Reaction was started by addition of 4 μl 5X RT Buffer [250 mM Tris-HCl, pH 8.3; 375 mM KCl; 15 mM MgCl_2_], 1 μl 0.1 M DTT, 20 U SUPERase^•^ In^™^ RNase Inhibitor, and 200 U TGIRT^™^-III Enzyme. Sample was incubated at 50°C for 5 minutes, followed by 2 hours at 57°C. Reaction was stopped by addition of 1 μl NaOH 5M. Sample was then incubated at 95°C for 3 minutes to detach the reverse transcriptase and degrade the RNA. cDNA cleanup was performed using Zymo RNA Clean & Concentrator-5 columns, following manufacturer instructions. PCR amplification of target intronic regions was performed using 10% of the purified cDNA for each primers pair and AccuPrime^™^ Taq DNA Polymerase (high fidelity), following manufacturer instructions. PCR products were resolved on a 2% Agarose gel, the bands of the expected size were gel-purified and pooled in equimolar amounts. Sample were then sheared by 20 cycles of sonication using a Diagenode Bioruptor^®^ sonicator [30’’ ON; 30’’ OFF; Power: high]. Libraries were prepared using the NEBNext^®^ Ultra^™^ II DNA Library Prep Kit for Illumina^®^, following manufacturer instructions.

#### DMS-MaPseq data analysis

Data analysis was performed using the RNA Framework v2.5 (http://www.rnaframework.com) (Incarnato, Neri, Anselmi, & Oliviero, 2016) All tools were used with default parameters, except for the *rf-map* tool that was called with the parameter -*mp “--very-sensitive-local”*. The percentage of mutations per base was calculated by normalizing the number of mutations on that base by the frequency of that base in the reference (e.g. the number of mutations occurring on A residues, divided by the total number of As present in the analyzed introns).

### Accession Numbers

RNA Sequencing and eCLIP data have been deposited in the NCBI GEO repository, accession number GSE113786. Data for circRNA in prostate cancer patients was obtained from Ref (S. Chen et al., 2019), NCBI GEO repository accession number: GSE113120.

## SUPPLEMENTARY FIGURE LEGENDS

**Figure S1.**
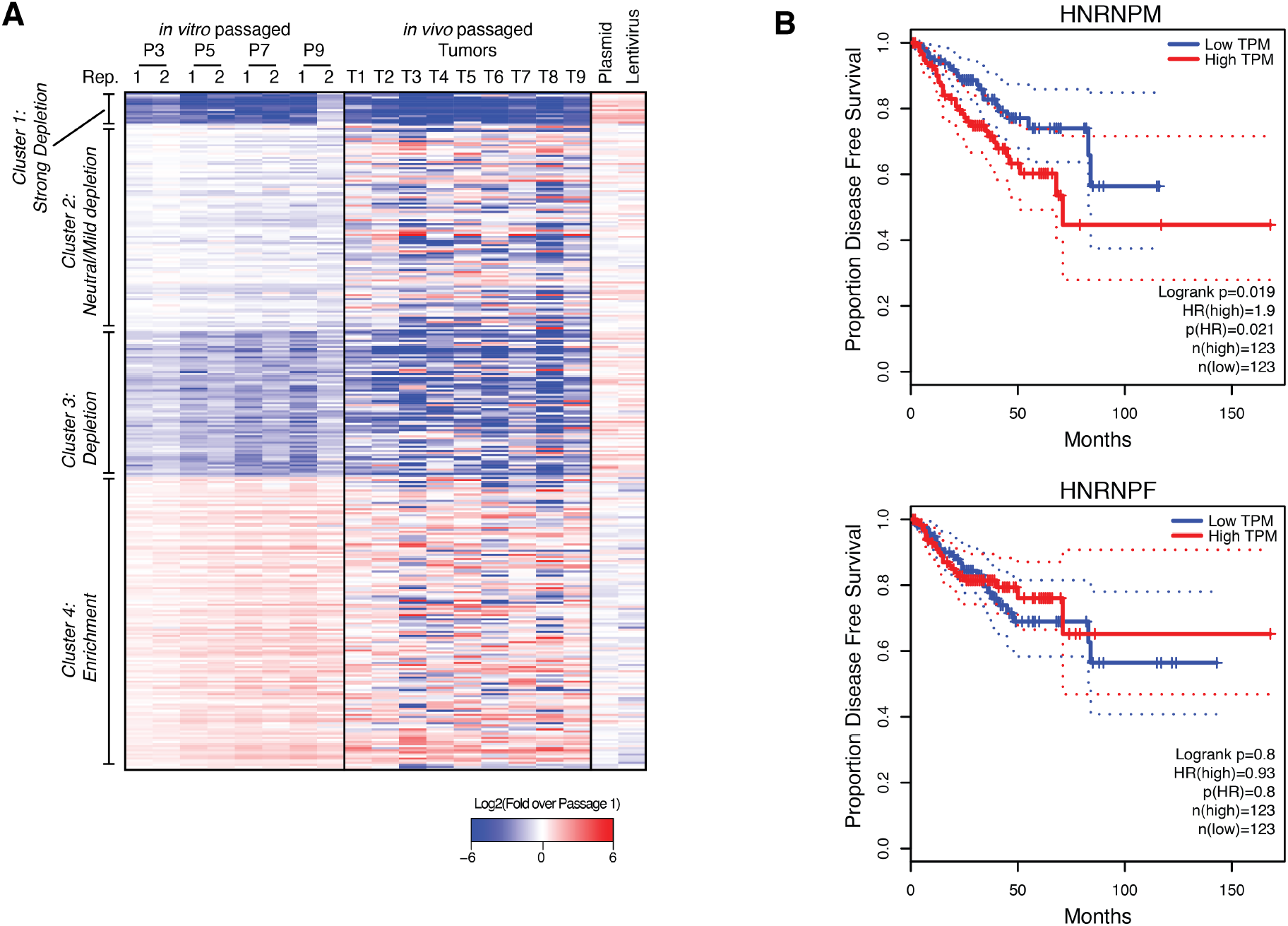
(Related to Figure 1): Verification of top hits in the pooled shRNA screen. **(A)** Heatmap showing overall hairpin enrichment over time in *in vitro* passaged LNCAP cells (left), *in vivo* LNCAP xenografts (center) as well as input PLKO.1 plasmid and lentiviral pools (right). Colors are shown in relation to the first passage of cells collected after completion of puromycin selection (P1), where blue indicates depletion and red indicates enrichment of the individual hairpin over time. **(B)** Disease-free survival curves of Prostate cancer patients that express high (>75^th^ percentile) or low (≤25^th^ percentile) of HNRNPM or HNRNPF. Data were plotted using GEPIA (Tang et al., 2017) and based on publically available TCGA datasets.

**Figure S2.**
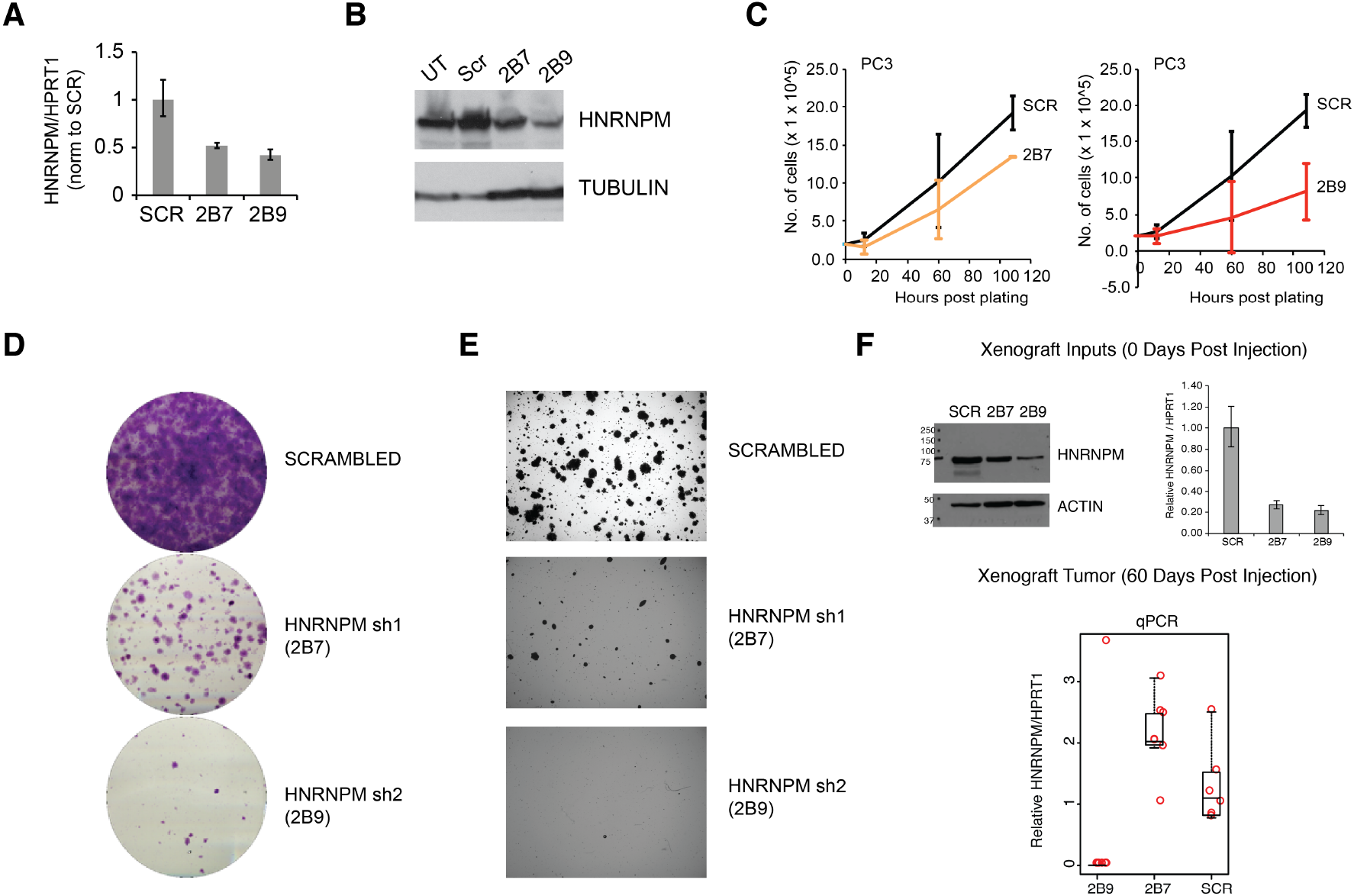
(Related to Figure 2): Impact of HNRNPM deficiency in an independent PCa cell line. HNRNPM RNA **(A)** and protein levels **(B)** upon expression of scrambled or HNRNPM specific shRNAs in PC3 cells. **(C)** Cell proliferation assays of PC3 cells expressing with either scrambled or HNRNPM specific shRNAs (2B7 and 2B9). **(D)** colony formation assays and **(E)** anchorage independent growth (soft agar assays)of shRNA treated PC3 cells **(F)** Western blotting and RNA analysis of cells used for xenograft experiments (top panel). qPCR analysis of RNA isolated from tumors harvested from xenografted mice at 60 days post injection (bottom panel). Each dot indicates one tumor.

**Figure S3.**
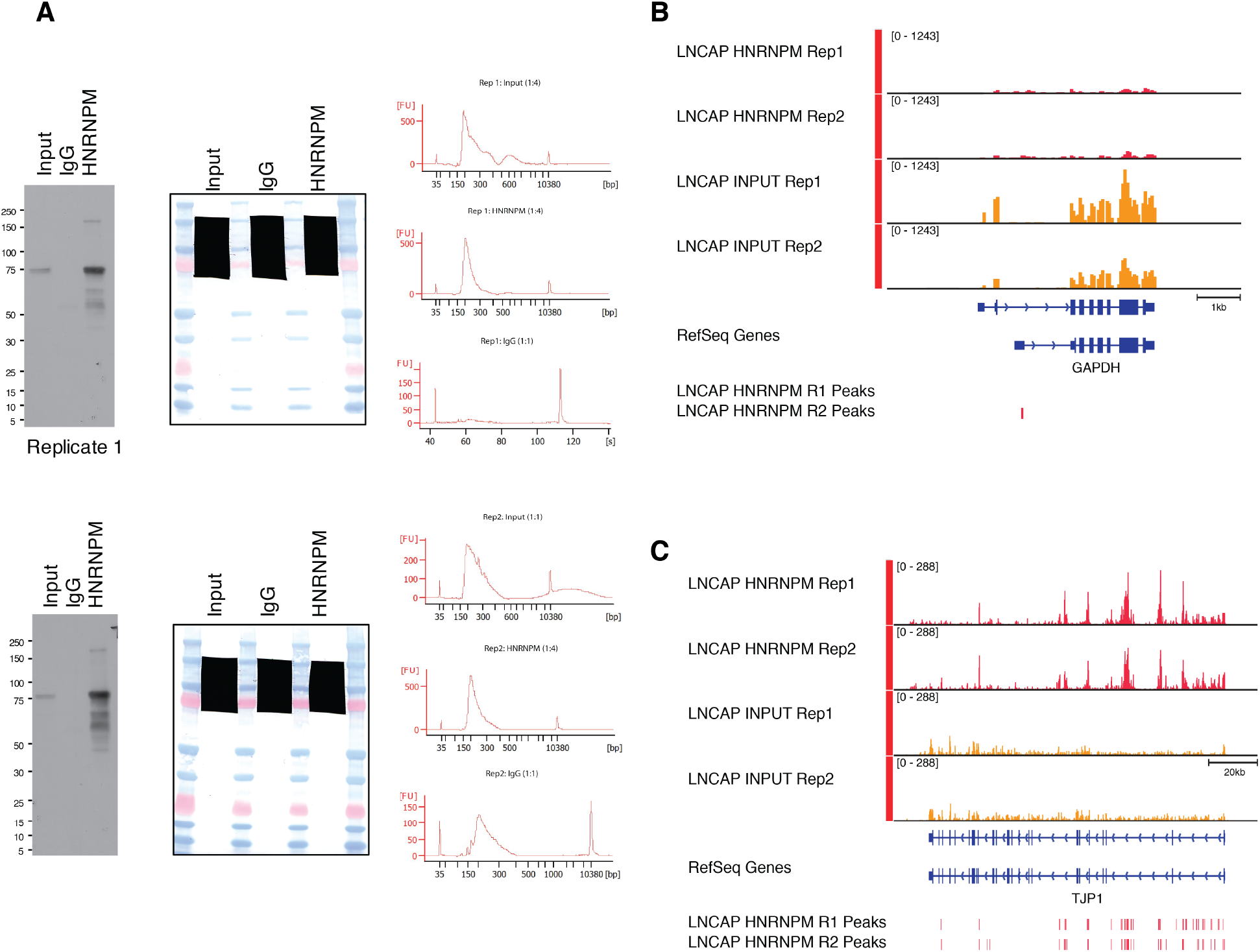
(related to Figure 3). eCLIP analysis of HNRNPM binding sites. **(A, left)** Immunoprecipitation-western blot validation of anti-HNRNPM antibody pull down efficiency in UV-crosslinked LNCAP lysates. **(A, center)** Scan of membrane showing regions excised for eCLIP library preparation. **(A, right)** High-sensitivity DNA Bioanalyzer traces of eCLIP libraries used for sequencing. **(B and C)** Read density of HNRNPM eCLIP at a HNRNPM bound locus, TJP1 **(C)** and a non-bound locus, GAPDH **(B)**. Bars indicate high confidence (p<0.05, fold enrichment > 2) called eCLIP peaks.

**Figure S4.**
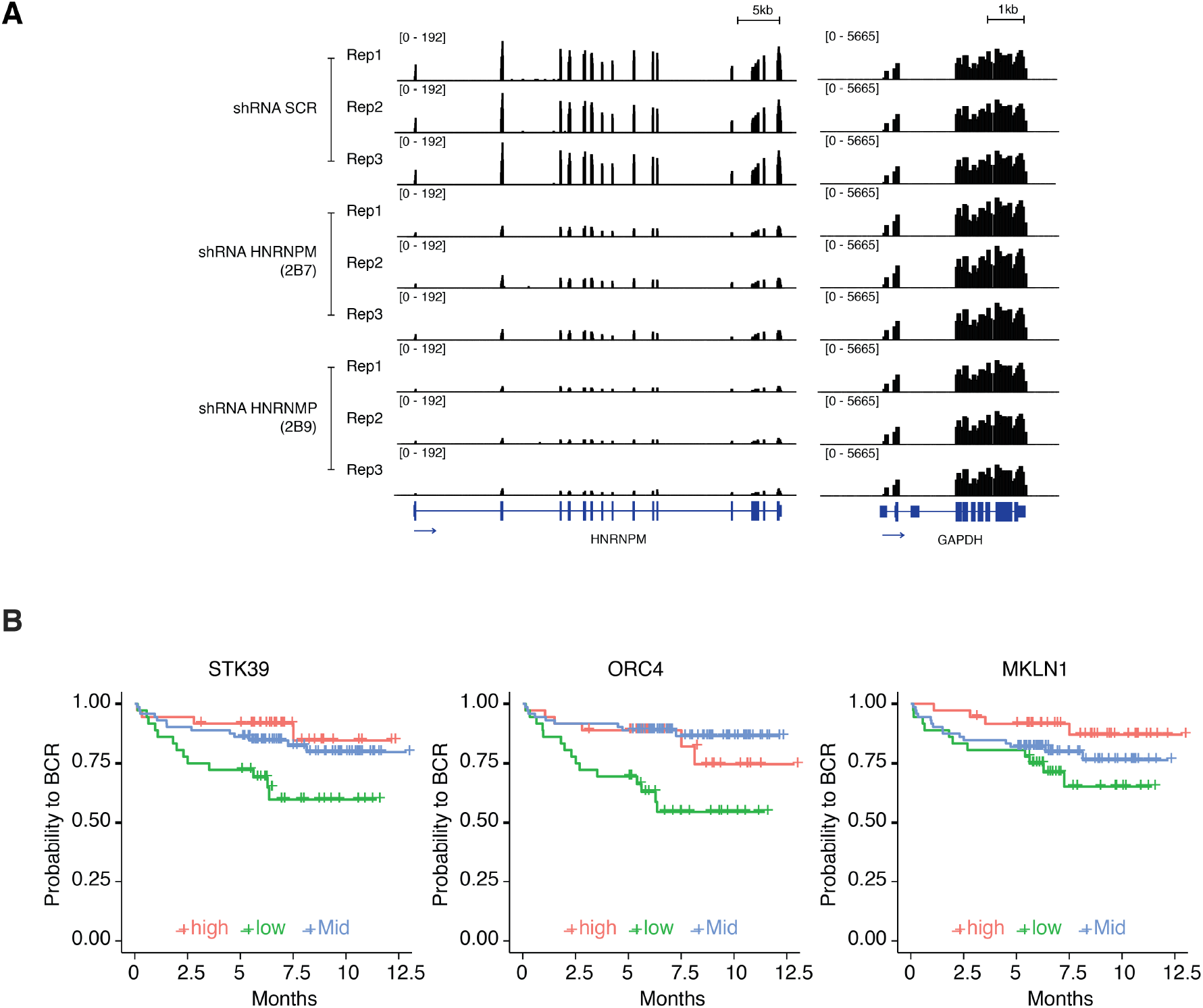
(related to Figure 4 and 5). Controls related to Figure 4 and 5. **(A)** RNA sequencing read density at the HNRNPM locus in scrambled and HNRNPM (2B7 and 2B9) shRNA treated cells. RNA sequencing tracks at the GAPDH housekeeping gene is shown as a control. **(B)** Biochemical relapse survival curves of prostate cancer patients expressing high (>75^th^ percentile), low (≤25^th^ percentile) or mid (25^th^-75^th^ percentile) levels of the indicated circular RNAs.

**Figure S5:**
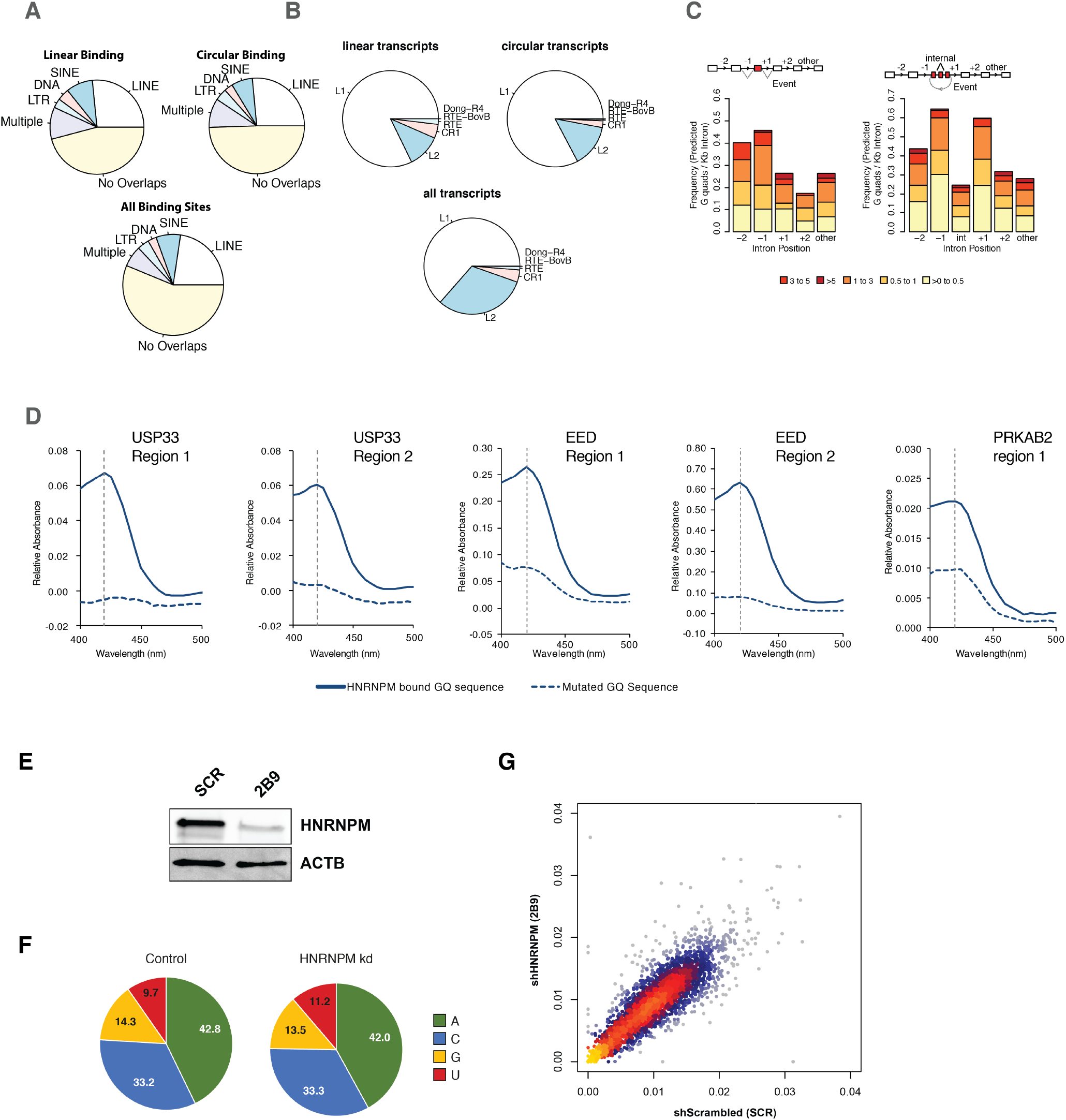
Structural Characterization of HNRNPM bound introns. **(A)** Distribution of unique HNRNPM peaks found in the immediate flanking upstream and downstream introns of mis-spliced linear (top left panel) or circular (top right panel) mis-splicing events that share a >10% overlap with one or multiple classes of 4 types of repetitive elements (LINEs, SINEs, DNA and LTR). The overlap of these elements with HNRNPM peaks found across the transcriptome is indicated in the bottom panel. **(B)** Types of LINE elements associated with HNRNPM binding sites at mis-spliced linear (top left panel) and circular (top right panel) transcripts, compared to that in regions bound by HNRNPM (bottom panel) across all transcripts **(C)** Distribution of G-quadruplex forming regions in the upstream (−1 or −2) or downstream (+1 or +2) introns relative to mis-spliced linear (left) or circular (right) events. Counts are normalized to the length of the intron. **(D)** Colorimetric, hemin based assay for RNA G-quadruplex folding. RNA oligos of HNRNPM binding sites of the USP33, EED and PRKAB2 genes were folded in the presence of potassium, and incubated with hemin. The presence of RNA G-quadruplexes in conjunction with hemin results in a peroxidase activity, which can be detected as an increase in absorbance at 420nm. As a negative control, the same test oligos with mutated GQ sites was used (dotted lines) **(E)** HNRNPM protein levels in cells used for targeted DMS-MaPseq. **(F)** Base mutation frequencies in Scrambled shRNA or HNRNPM (2B9) shRNA treated LNCAP cells. **(G)** Overall correlation of the structure probing data for Scrambled shRNA and HNRNPM shRNA (2B9) treated cells. Representative HNRNPM bound sites in the introns flanking mis-splicing events for PRKAB2, USP33, ZNF548, ZNF304, GMPR2, GAPDH were used for this analysis.

**Figure S6:**
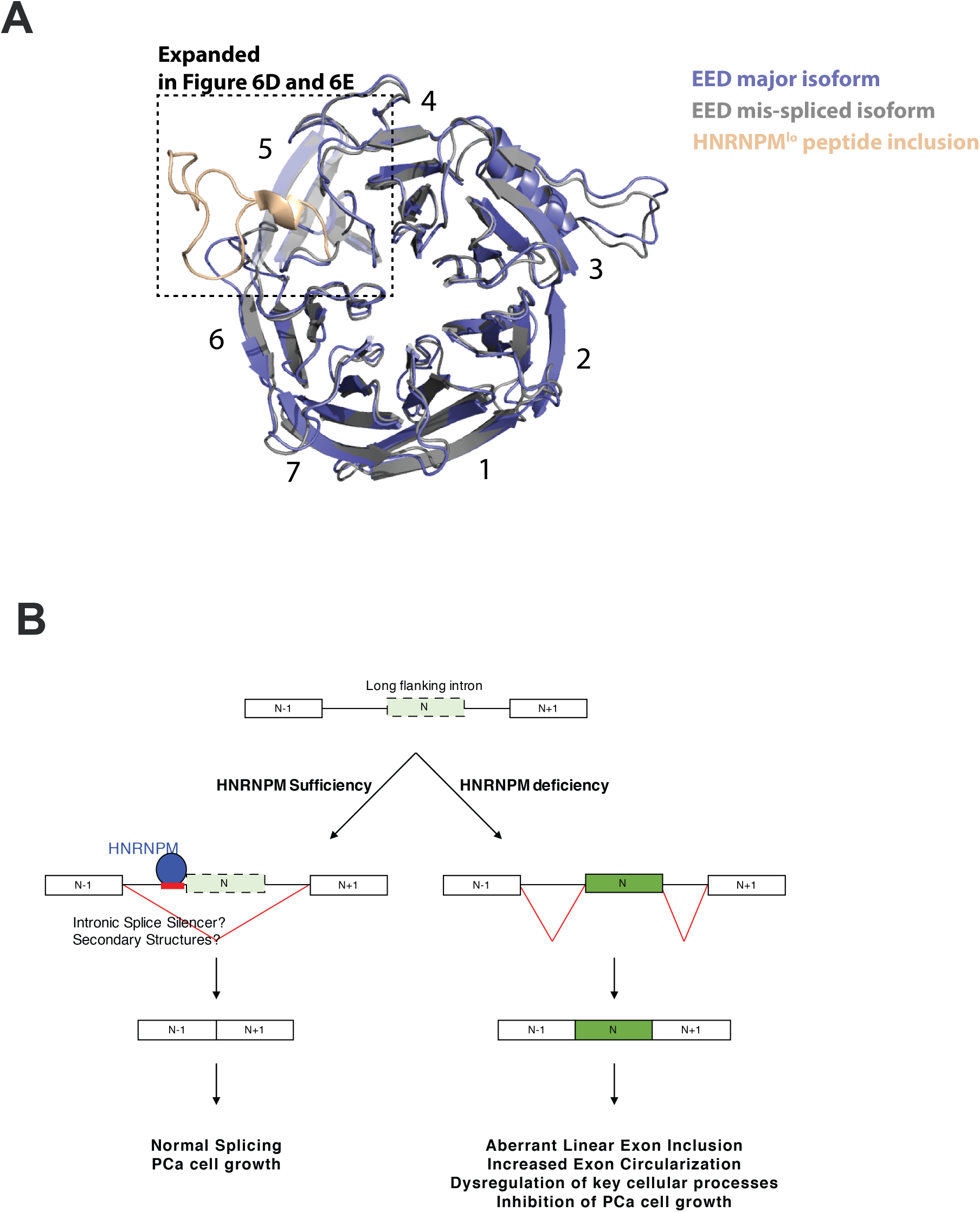
HNRNPM regulates a multigenic splicing program to maintain cell proliferation. **(A)** Structure of the HNRNPM regulated WD40 domain in wildtype EED protein (Grey), superimposed on the predicted structure of the HNRNPM-dependent EED isoform (blue) that contains the 25 aa long peptide coded by the exon regulated by HNRNPM. The new peptide generated by the splicing event is depicted in orange. WD domains in EED are numbered. **(B)** Model summarizing the role of HNRNPM in PCa cells.

## SUPPLEMENTARY TABLE LEGENDS

**Table S1:** Table of shRNAs and sequences used for pooled screens (Related to Figure 1 and S1)

**Table S2:** Rotational Gene Set Analysis (ROAST) results: Tumor versus Input (P1) (Related to Figure 1C)

**Table S3**: Rotational Gene Set Analysis (ROAST) results: In vitro (Related to Figure 1C)

**Table S4:** Gene Expression Changes in all HNRNPM bound genes (Related to Figure 4A)

**Table S5:** Binding Status of all significantly differentially expressed genes in HNRNPM knockdown (Related to Figure 4A)

**Table S6**: Binding Status of Significantly Changed Skipped Exon Events (Related to Figure 4B and 4C)

**Table S7:** Binding Status of Significantly Changed Circular RNAs (Related to Figure 5A)

**Table S8**: Sequences used for Bichromatic Splicing Reporter (Related to Figure 6)

**Table S9**: Sequences of RNA oligos used for RNA G quadruplex Hemin Assays (Related to Figure S5D)

## REFERENCES

Aktas, T., Avsar Ilik, I., Maticzka, D., Bhardwaj, V., Pessoa Rodrigues, C., Mittler, G.,… Akhtar, A. (2017). DHX9 suppresses RNA processing defects originating from the Alu invasion of the human genome. Nature, 544(7648), 115–119. doi:10.1038/nature21715

Baralle, F. E., & Giudice, J. (2017). Alternative splicing as a regulator of development and tissue identity. Nat Rev Mol Cell Biol, 18(7), 437–451. doi:10.1038/nrm.2017.27

Bartram, M. P., Mishra, T., Reintjes, N., Fabretti, F., Gharbi, H., Adam, A. C.,… Muller, R. U. (2017). Characterization of a splice-site mutation in the tumor suppressor gene FLCN associated with renal cancer. BMC Med Genet, 18(1), 53. doi:10.1186/s12881-017-0416-5

Blanchette, M., & Chabot, B. (1999). Modulation of exon skipping by high-affinity hnRNP A1-binding sites and by intron elements that repress splice site utilization. EMBO J, 18(7), 1939–1952. doi:10.1093/emboj/18.7.1939

Brannan, K. W., Jin, W., Huelga, S. C., Banks, C. A., Gilmore, J. M., Florens, L.,… Yeo, G. W. (2016). SONAR Discovers RNA-Binding Proteins from Analysis of Large-Scale Protein-Protein Interactomes. Mol Cell, 64(2), 282–293. doi:10.1016/j.molcel.2016.09.003

Braun, C. J., Stanciu, M., Boutz, P. L., Patterson, J. C., Calligaris, D., Higuchi, F.,… Lees, J. A. (2017). Coordinated Splicing of Regulatory Detained Introns within Oncogenic Transcripts Creates an Exploitable Vulnerability in Malignant Glioma. Cancer Cell, 32(4), 411–426 e411. doi:10.1016/j.ccell.2017.08.018

Chen, L., Chen, J. Y., Huang, Y. J., Gu, Y., Qiu, J., Qian, H.,…. Fu, X. D. (2018). The Augmented R-Loop Is a Unifying Mechanism for Myelodysplastic Syndromes Induced by High-Risk Splicing Factor Mutations. Mol Cell, 69(3), 412–425 e416. doi:10.1016/j.molcel.2017.12.029

Chen, S., Huang, V., Xu, X., Livingstone, J., Soares, F., Jeon, J.,… He, H. H. (2019). Widespread and Functional RNA Circularization in Localized Prostate Cancer. Cell, 176(4), 831–843 e822. doi:10.1016/j.cell.2019.01.025

Damianov, A., Ying, Y., Lin, C. H., Lee, J. A., Tran, D., Vashisht, A. A.,… Black, D. L. (2016). Rbfox Proteins Regulate Splicing as Part of a Large Multiprotein Complex LASR. Cell, 165(3), 606–619. doi:10.1016/j.cell.2016.03.040

Dewaele, M., Tabaglio, T., Willekens, K., Bezzi, M., Teo, S. X., Low, D. H.,… Guccione, E. (2016). Antisense oligonucleotide-mediated MDM4 exon 6 skipping impairs tumor growth. J Clin Invest, 126(1), 68–84. doi:10.1172/JCI82534

Dobin, A., Davis, C. A., Schlesinger, F., Drenkow, J., Zaleski, C., Jha, S.,… Gingeras, T. R. (2013). STAR: ultrafast universal RNA-seq aligner. Bioinformatics, 29(1), 15–21. doi:10.1093/bioinformatics/bts635

Ellis, J. D., Barrios-Rodiles, M., Colak, R., Irimia, M., Kim, T., Calarco, J. A.,… Blencowe, B. J. (2012). Tissue-specific alternative splicing remodels protein-protein interaction networks. Mol Cell, 46(6), 884–892. doi:10.1016/j.molcel.2012.05.037

Fagoonee, S., Picco, G., Orso, F., Arrigoni, A., Longo, D. L., Forni, M.,… Altruda, F. (2017). The RNA-binding protein ESRP1 promotes human colorectal cancer progression. Oncotarget, 8(6), 10007–10024. doi:10.18632/oncotarget.14318

Gargiulo, G., Serresi, M., Cesaroni, M., Hulsman, D., & van Lohuizen, M. (2014). In vivo shRNA screens in solid tumors. Nat Protoc, 9(12), 2880–2902. doi:10.1038/nprot.2014.185

Graubert, T. A., Shen, D., Ding, L., Okeyo-Owuor, T., Lunn, C. L., Shao, J.,… Walter, M. J. (2011). Recurrent mutations in the U2AF1 splicing factor in myelodysplastic syndromes. Nat Genet, 44(1), 53–57. doi:10.1038/ng.1031

Hang, X., Li, P., Li, Z., Qu, W., Yu, Y., Li, H.,… Zhang, C. (2009). Transcription and splicing regulation in human umbilical vein endothelial cells under hypoxic stress conditions by exon array. BMC Genomics, 10, 126. doi:10.1186/1471-2164-10-126

Hansen, T. B., Jensen, T. I., Clausen, B. H., Bramsen, J. B., Finsen, B., Damgaard, C. K., & Kjems, J. (2013). Natural RNA circles function as efficient microRNA sponges. Nature, 495(7441), 384–388. doi:10.1038/nature11993

Haque, N., Ouda, R., Chen, C., Ozato, K., & Hogg, J. R. (2018). ZFR coordinates crosstalk between RNA decay and transcription in innate immunity. Nat Commun, 9(1), 1145. doi:10.1038/s41467-018-03326-5

Hsu, T. Y., Simon, L. M., Neill, N. J., Marcotte, R., Sayad, A., Bland, C. S.,… Westbrook, T. F. (2015). The spliceosome is a therapeutic vulnerability in MYC-driven cancer. Nature, 525(7569), 384–388. doi:10.1038/nature14985

Huang, H., Zhang, J., Harvey, S. E., Hu, X., & Cheng, C. (2017). RNA G-quadruplex secondary structure promotes alternative splicing via the RNA-binding protein hnRNPF. Genes Dev, 31(22), 2296–2309. doi:10.1101/gad.305862.117

Huelga, S. C., Vu, A. Q., Arnold, J. D., Liang, T. Y., Liu, P. P., Yan, B. Y.,… Yeo, G. W. (2012). Integrative genome-wide analysis reveals cooperative regulation of alternative splicing by hnRNP proteins. Cell Rep, 1(2), 167–178. doi:10.1016/j.celrep.2012.02.001

Incarnato, D., Neri, F., Anselmi, F., & Oliviero, S. (2016). RNA structure framework: automated transcriptome-wide reconstruction of RNA secondary structures from high-throughput structure probing data. Bioinformatics, 32(3), 459–461. doi:10.1093/bioinformatics/btv571

Jeck, W. R., Sorrentino, J. A., Wang, K., Slevin, M. K., Burd, C. E., Liu, J.,… Sharpless, N. E. (2013). Circular RNAs are abundant, conserved, and associated with ALU repeats. RNA, 19(2), 141–157. doi:10.1261/rna.035667.112

Jung, H., Lee, D., Lee, J., Park, D., Kim, Y. J., Park, W. Y.,… Lee, E. (2015). Intron retention is a widespread mechanism of tumor-suppressor inactivation. Nat Genet, 47(11), 1242–1248. doi:10.1038/ng.3414

Kelley, D. R., Hendrickson, D. G., Tenen, D., & Rinn, J. L. (2014). Transposable elements modulate human RNA abundance and splicing via specific RNA-protein interactions. Genome Biol, 15(12), 537. doi:10.1186/s13059-014-0537-5

King, I. F., Yandava, C. N., Mabb, A. M., Hsiao, J. S., Huang, H. S., Pearson, B. L.,… Zylka, M. J. (2013). Topoisomerases facilitate transcription of long genes linked to autism. Nature, 501(7465), 58–62. doi:10.1038/nature12504

Koh, C. M., Bezzi, M., Low, D. H., Ang, W. X., Teo, S. X., Gay, F. P.,… Guccione, E. (2015). MYC regulates the core pre-mRNA splicing machinery as an essential step in lymphomagenesis. Nature, 523(7558), 96–100. doi:10.1038/nature14351

Lareau, L. F., & Brenner, S. E. (2015). Regulation of splicing factors by alternative splicing and NMD is conserved between kingdoms yet evolutionarily flexible. Mol Biol Evol, 32(4), 1072–1079. doi:10.1093/molbev/msv002

Li, H., Liu, J., Fang, Y., Qin, Y., Xu, S., Liu, Y., & Wang, E. (2013). G-quadruplex-based ultrasensitive and selective detection of histidine and cysteine. Biosens Bioelectron, 41, 563–568. doi:10.1016/j.bios.2012.09.024

Liu, L. L., Xie, N., Sun, S., Plymate, S., Mostaghel, E., & Dong, X. (2014). Mechanisms of the androgen receptor splicing in prostate cancer cells. Oncogene, 33(24), 3140–3150. doi:10.1038/onc.2013.284

Low, D. (2017). SPLINTER: Splice Interpreter Of Transcripts. version 1.4.0.

Makino, Y., Kanopka, A., Wilson, W. J., Tanaka, H., & Poellinger, L. (2002). Inhibitory PAS domain protein (IPAS) is a hypoxia-inducible splicing variant of the hypoxia-inducible factor-3alpha locus. J Biol Chem, 277(36), 32405–32408. doi:10.1074/jbc.C200328200

Martinez-Contreras, R., Fisette, J. F., Nasim, F. U., Madden, R., Cordeau, M., & Chabot, B. (2006). Intronic binding sites for hnRNP A/B and hnRNP F/H proteins stimulate pre-mRNA splicing. PLoS Biol, 4(2), e21. doi:10.1371/journal.pbio.0040021

Matlin, A. J., Clark, F., & Smith, C. W. (2005). Understanding alternative splicing: towards a cellular code. Nat Rev Mol Cell Biol, 6(5), 386–398. doi:10.1038/nrm1645

Mayeda, A., Helfman, D. M., & Krainer, A. R. (1993). Modulation of exon skipping and inclusion by heterogeneous nuclear ribonucleoprotein A1 and pre-mRNA splicing factor SF2/ASF. Mol Cell Biol, 13(5), 2993–3001.

Mayeda, A., & Krainer, A. R. (1992). Regulation of alternative pre-mRNA splicing by hnRNP A1 and splicing factor SF2. Cell, 68(2), 365–375.

Memczak, S., Jens, M., Elefsinioti, A., Torti, F., Krueger, J., Rybak, A.,… Rajewsky, N. (2013). Circular RNAs are a large class of animal RNAs with regulatory potency. Nature, 495(7441), 333–338. doi:10.1038/nature11928

Naro, C., Jolly, A., Di Persio, S., Bielli, P., Setterblad, N., Alberdi, A. J.,… Sette, C. (2017). An Orchestrated Intron Retention Program in Meiosis Controls Timely Usage of Transcripts during Germ Cell Differentiation. Dev Cell, 41(1), 82–93 e84. doi:10.1016/j.devcel.2017.03.003

Orengo, J. P., Bundman, D., & Cooper, T. A. (2006). A bichromatic fluorescent reporter for cell-based screens of alternative splicing. Nucleic Acids Res, 34(22), e148. doi:10.1093/nar/gkl967

Papaemmanuil, E., Cazzola, M., Boultwood, J., Malcovati, L., Vyas, P., Bowen, D.,… Chronic Myeloid Disorders Working Group of the International Cancer Genome, C. (2011). Somatic SF3B1 mutation in myelodysplasia with ring sideroblasts. N Engl J Med, 365(15), 1384–1395. doi:10.1056/NEJMoa1103283

Pimentel, H., Parra, M., Gee, S. L., Mohandas, N., Pachter, L., & Conboy, J. G. (2016). A dynamic intron retention program enriched in RNA processing genes regulates gene expression during terminal erythropoiesis. Nucleic Acids Res, 44(2), 838–851. doi:10.1093/nar/gkv1168

Puente, X. S., Bea, S., Valdes-Mas, R., Villamor, N., Gutierrez-Abril, J., Martin-Subero, J. I.,… Campo, E. (2015). Non-coding recurrent mutations in chronic lymphocytic leukaemia. Nature, 526(7574), 519–524. doi:10.1038/nature14666

Rogic, S., Montpetit, B., Hoos, H. H., Mackworth, A. K., Ouellette, B. F., & Hieter, P. (2008). Correlation between the secondary structure of pre-mRNA introns and the efficiency of splicing in Saccharomyces cerevisiae. BMC Genomics, 9, 355. doi:10.1186/1471-2164-9-355

Sahakyan, A. B., & Balasubramanian, S. (2016). Long genes and genes with multiple splice variants are enriched in pathways linked to cancer and other multigenic diseases. BMC Genomics, 17, 225. doi:10.1186/s12864-016-2582-9

Schwerk, C., & Schulze-Osthoff, K. (2005). Regulation of apoptosis by alternative pre-mRNA splicing. Mol Cell, 19(1), 1–13. doi:10.1016/j.molcel.2005.05.026

Sebestyen, E., Zawisza, M., & Eyras, E. (2015). Detection of recurrent alternative splicing switches in tumor samples reveals novel signatures of cancer. Nucleic Acids Res, 43(3), 1345–1356. doi:10.1093/nar/gku1392

Serresi, M., Gargiulo, G., Proost, N., Siteur, B., Cesaroni, M., Koppens, M.,… van Lohuizen, M. (2016). Polycomb Repressive Complex 2 Is a Barrier to KRAS-Driven Inflammation and Epithelial-Mesenchymal Transition in Non-Small-Cell Lung Cancer. Cancer Cell, 29(1), 17–31. doi:10.1016/j.ccell.2015.12.006

Shalgi, R., Hurt, J. A., Lindquist, S., & Burge, C. B. (2014). Widespread inhibition of posttranscriptional splicing shapes the cellular transcriptome following heat shock. Cell Rep, 7(5), 1362–1370. doi:10.1016/j.celrep.2014.04.044

Shapiro, I. M., Cheng, A. W., Flytzanis, N. C., Balsamo, M., Condeelis, J. S., Oktay, M. H.,… Gertler, F. B. (2011). An EMT-driven alternative splicing program occurs in human breast cancer and modulates cellular phenotype. PLoS Genet, 7(8), e1002218. doi:10.1371/journal.pgen.1002218

Shen, S., Park, J. W., Lu, Z. X., Lin, L., Henry, M. D., Wu, Y. N.,… Xing, Y. (2014). rMATS: robust and flexible detection of differential alternative splicing from replicate RNA-Seq data. Proc Natl Acad Sci U S A, 111(51), E5593–5601. doi:10.1073/pnas.1419161111

Shepard, S., McCreary, M., & Fedorov, A. (2009). The peculiarities of large intron splicing in animals. PLoS One, 4(11), e7853. doi:10.1371/journal.pone.0007853

Shi, J., Wang, E., Zuber, J., Rappaport, A., Taylor, M., Johns, C.,… Vakoc, C. R. (2013). The Polycomb complex PRC2 supports aberrant self-renewal in a mouse model of MLL-AF9;Nras(G12D) acute myeloid leukemia. Oncogene, 32(7), 930–938. doi:10.1038/onc.2012.110

Solnick, D., & Lee, S. I. (1987). Amount of RNA secondary structure required to induce an alternative splice. Mol Cell Biol, 7(9), 3194–3198.

t Hoen, P. A., Hirsch, M., de Meijer, E. J., de Menezes, R. X., van Ommen, G. J., & den Dunnen, J. T. (2011). mRNA degradation controls differentiation state-dependent differences in transcript and splice variant abundance. Nucleic Acids Res, 39(2), 556–566. doi:10.1093/nar/gkq790

Tang, Z., Li, C., Kang, B., Gao, G., Li, C., & Zhang, Z. (2017). GEPIA: a web server for cancer and normal gene expression profiling and interactive analyses. Nucleic Acids Res, 45(W1), W98–W102. doi:10.1093/nar/gkx247

Todaro, M., Gaggianesi, M., Catalano, V., Benfante, A., Iovino, F., Biffoni, M.,… Stassi, G. (2014). CD44v6 is a marker of constitutive and reprogrammed cancer stem cells driving colon cancer metastasis. Cell Stem Cell, 14(3), 342–356. doi:10.1016/j.stem.2014.01.009

Trapnell, C., Roberts, A., Goff, L., Pertea, G., Kim, D., Kelley, D. R.,… Pachter, L. (2012). Differential gene and transcript expression analysis of RNA-seq experiments with TopHat and Cufflinks. Nat Protoc, 7(3), 562–578. doi:10.1038/nprot.2012.016

Ueda, J., Matsuda, Y., Yamahatsu, K., Uchida, E., Naito, Z., Korc, M., & Ishiwata, T. (2014). Epithelial splicing regulatory protein 1 is a favorable prognostic factor in pancreatic cancer that attenuates pancreatic metastases. Oncogene, 33(36), 4485–4495. doi:10.1038/onc.2013.392

van Bokhoven, A., Varella-Garcia, M., Korch, C., Johannes, W. U., Smith, E. E., Miller, H. L.,… Lucia, M. S. (2003). Molecular characterization of human prostate carcinoma cell lines. Prostate, 57(3), 205–225. doi:10.1002/pros.10290

Van Nostrand, E. L., Pratt, G. A., Shishkin, A. A., Gelboin-Burkhart, C., Fang, M. Y., Sundararaman, B.,… Yeo, G. W. (2016). Robust transcriptome-wide discovery of RNA-binding protein binding sites with enhanced CLIP (eCLIP). Nat Methods, 13(6), 508–514. doi:10.1038/nmeth.3810

Venter, J. C., Adams, M. D., Myers, E. W., Li, P. W., Mural, R. J., Sutton, G. G.,… Zhu, X. (2001). The sequence of the human genome. Science, 291(5507), 1304–1351. doi:10.1126/science.1058040

Wahl, M. C., Will, C. L., & Luhrmann, R. (2009). The spliceosome: design principles of a dynamic RNP machine. Cell, 136(4), 701–718. doi:10.1016/j.cell.2009.02.009

Wang, L., Lawrence, M. S., Wan, Y., Stojanov, P., Sougnez, C., Stevenson, K.,… Wu, C. J. (2011). SF3B1 and other novel cancer genes in chronic lymphocytic leukemia. N Engl J Med, 365(26), 2497–2506. doi:10.1056/NEJMoa1109016

Wu, D., Lim, E., Vaillant, F., Asselin-Labat, M. L., Visvader, J. E., & Smyth, G. K. (2010). ROAST: rotation gene set tests for complex microarray experiments. Bioinformatics, 26(17), 2176–2182. doi:10.1093/bioinformatics/btq401

Xu, Y., Gao, X. D., Lee, J. H., Huang, H., Tan, H., Ahn, J.,… Cheng, C. (2014). Cell type-restricted activity of hnRNPM promotes breast cancer metastasis via regulating alternative splicing. Genes Dev, 28(11), 1191–1203. doi:10.1101/gad.241968.114

Yang, X., Coulombe-Huntington, J., Kang, S., Sheynkman, G. M., Hao, T., Richardson, A.,… Vidal, M. (2016). Widespread Expansion of Protein Interaction Capabilities by Alternative Splicing. Cell, 164(4), 805–817. doi:10.1016/j.cell.2016.01.029

Yoshida, K., Sanada, M., Shiraishi, Y., Nowak, D., Nagata, Y., Yamamoto, R.,… Ogawa, S. (2011). Frequent pathway mutations of splicing machinery in myelodysplasia. Nature, 478(7367), 64–69. doi:10.1038/nature10496

Zhang, X. O., Wang, H. B., Zhang, Y., Lu, X., Chen, L. L., & Yang, L. (2014). Complementary sequence-mediated exon circularization. Cell, 159(1), 134–147. doi:10.1016/j.cell.2014.09.001

Zheng, S., Vuong, B. Q., Vaidyanathan, B., Lin, J. Y., Huang, F. T., & Chaudhuri, J. (2015). Non-coding RNA Generated following Lariat Debranching Mediates Targeting of AID to DNA. Cell, 161(4), 762–773. doi:10.1016/j.cell.2015.03.020

Zhu, J., Mayeda, A., & Krainer, A. R. (2001). Exon identity established through differential antagonism between exonic splicing silencer-bound hnRNP A1 and enhancer-bound SR proteins. Mol Cell, 8(6), 1351–1361.

Zubradt, M., Gupta, P., Persad, S., Lambowitz, A. M., Weissman, J. S., & Rouskin, S. (2017). DMS-MaPseq for genome-wide or targeted RNA structure probing in vivo. Nat Methods, 14(1), 75–82. doi:10.1038/nmeth.4057

